# Gut microbiota and fecal short chain fatty acids differ with adiposity and country of origin: The METS-Microbiome Study

**DOI:** 10.1101/2023.03.21.533195

**Authors:** Gertrude Ecklu-Mensah, Candice Choo-Kang, Maria Gjerstad Maseng, Sonya Donato, Pascal Bovet, Kweku Bedu-Addo, Jacob Plange-Rhule, Terrence E. Forrester, Estelle V. Lambert, Dale Rae, Amy Luke, Brian T. Layden, Stephen O’Keefe, Jack A. Gilbert, Lara R. Dugas

## Abstract

The relationship between the gut microbiota, short chain fatty acid (SCFA) metabolism, and obesity remains unclear due to conflicting reports from studies with limited statistical power. Additionally, this association has rarely been explored in large scale diverse populations. Here, we investigated associations between fecal microbial composition, predicted metabolic potential, SCFA concentrations, and obesity in a large (*N* = 1,934) adult cohort of African-origin spanning the epidemiologic transition, from Ghana, South Africa, Jamaica, Seychelles, and the United States (US). The greatest gut microbiota diversity and total fecal SCFA concentration was found in the Ghanaian population, while the lowest levels were found in the US population, respectively representing the lowest and the highest end of the epidemiologic transition spectrum. Country-specific bacterial taxa and predicted-functional pathways were observed, including an increased prevalence of *Prevotella*, *Butyrivibrio*, *Weisella* and *Romboutsia* in Ghana and South Africa, while *Bacteroides* and *Parabacteroides* were enriched in Jamaican and the US populations. Importantly, ’VANISH’ taxa, including *Butyricicoccus and Succinivibrio*, were significantly enriched in the Ghanaian cohort, reflecting the participants’ traditional lifestyles. Obesity was significantly associated with lower SCFA concentrations, a decrease in microbial richness, and dissimilarities in community composition, and reduction in the proportion of SCFA synthesizing bacteria including *Oscillospira*, *Christensenella*, *Eubacterium*, *Alistipes*, *Clostridium* and *Odoribacter*. Further, the predicted proportions of genes in the lipopolysaccharide (LPS) synthesis pathway were enriched in obese individuals, while genes associated with butyrate synthesis via the dominant pyruvate pathway were significantly reduced in obese individuals. Using machine learning, we identified features predictive of metabolic state and country of origin. Country of origin could accurately be predicted by the fecal microbiota (AUC = 0.97), whereas obesity could not be predicted as accurately (AUC = 0.65). Participant sex (AUC = 0.75), diabetes status (AUC = 0.63), hypertensive status (AUC = 0.65), and glucose status (AUC = 0.66) could all be predicted with different success. Interestingly, within country, the predictive accuracy of the microbiota for obesity was inversely correlated to the epidemiological transition, being greatest in Ghana (AUC = 0.57). Collectively, our findings reveal profound variation in the gut microbiota, inferred functional pathways, and SCFA synthesis as a function of country of origin. While obesity could be predicted accurately from the microbiota, the variation in accuracy in parallel with the epidemiological transition suggests that differences in the microbiota between obesity and non-obesity may be larger in low-to-middle countries compared to high-income countries. Further examination of independent study populations using multi-omic approaches will be necessary to determine the factors that drive this association.

## Introduction

Obesity, which affects more than 600 million adults worldwide (“Obesity and Overweight” n.d.), over a third of Americans (Hales et al. 2020), and accounts for over 60% of deaths related to high body mass index (BMI) (Tseng and Wu 2019), remains an ongoing global health epidemic that continues to worsen at an alarming rate. A major driver of obesity is the adoption of a western lifestyle, which is characterized by excessive consumption of ultra-processed foods. Obesity is a major risk factor for type 2 diabetes, and according to the most recent National Diabetes Statistics Report almost 13% of the adult US population now have diabetes. Not only do 49.6% of adult African Americans present with obesity but over 17% of them now have diabetes, and are 1.5 times as likely to present with type 2 diabetes compared to whites (“National Diabetes Statistics Report” 2022). Populations of African-origin outside of the US are experiencing similar fates, as the prevalence of obesity among adults living in Sub-Saharan Africa is greater than 13%, and higher than the global obesity prevalence for adults (Agyemang et al. 2016). This has been accompanied by dramatic increases in the prevalence of non-communicable diseases such as type two diabetes and hypertension among people of African-origin (Roth et al. 2020; Gouda et al. 2019). Therefore, disrupting the rapidly expanding obesity epidemic, particularly among African-origin populations is critical to controlling the cardiometabolic disorder epidemic (Geng et al. 2022). However, successfully managing and treating obesity and its comorbidities, and specifically maintaining weight loss long-term, is particularly challenging due to an incomplete understanding of the heterogeneous and complex etiopathology, as well as additional challenges facing populations experiencing rapid urbanization (Nordmo, Danielsen, and Nordmo 2020; Geng et al. 2022; Barone et al. 2022). The epidemiologic transition is a model able to capture these shifts in dietary and rural to urban movements and is characterized by diets that are high in ultra-processed foods with a significant loss in fiber, as evidenced in the US, where less than 50% of the population meet dietary fiber recommendations (Dahl and Stewart 2015).

Gut microbial ecology and metabolism play pivotal roles in the onset and progression of obesity and its related metabolic disorders (Ley 2010). Obese and lean individuals have reported differences in the composition and functional potential of the gut microbiome, with an overall reduction in species diversity in the obese gut (Dugas, Bernabé, et al. 2018; Greenblum, Turnbaugh, and Borenstein 2012; Jumpertz et al. 2011; Ley et al. 2006; Turnbaugh et al. 2009; Le Chatelier et al. 2013), additionally, fecal microbiota transfer from obese donors to mouse models can recapitulate the obese phenotype (Turnbaugh et al. 2006, 2008; Ridaura et al. 2013). Further, fecal microbiota transplant from healthy donors into patients with obese and metabolic syndrome has been shown to improve markers of metabolic health in the recipients (Vrieze et al. 2012). While these studies suggest that modification of microbial ecology may offer new options for the treatment and prevention of obesity, the mechanism that drives the microbiota-obesity relationship is not fully understood. The microbiota may facilitate greater energy exploitation from food, and storage capacity by the host (Turnbaugh et al. 2006; DiBaise et al. 2008), influencing adipose tissue composition and fat mass gain, as well as providing chronic low-grade inflammation and insulin resistance (Cani and Delzenne 2009; J. L. Sonnenburg and Bäckhed 2016).

Among the numerous microbial metabolites modulating obesity, there is an ever-growing interest in the role of short-chain fatty acids (SCFAs), which includes butyrate, acetate, and propionate as potential biomarkers for metabolic health as well as therapeutic targets. SCFAs derive primarily from microbial fermentation of non-digestible dietary fiber in the colon. They have many effects on host metabolism including serving as an energy source for host colonocytes, used as precursors for the biosynthesis of cholesterol, lipids, proteins and regulating gut barrier activities (Dalile et al. 2019; Koh et al. 2016; van der Hee and Wells 2021). Human and animal studies demonstrate a protective role of SCFAs in obesity and metabolic disease. In experimental animal models, SCFA supplementation reduces body weight, improves insulin sensitivity, and reduces obesity-associated inflammation (Vinolo et al. 2011; Gao et al. 2009; Henagan et al. 2015; Lu et al. 2016; Bonomo et al. 2020). In humans, increased gut production of butyrate correlates with improved insulin response after an oral glucose-tolerance test (Sanna et al. 2019). Although increased SCFA levels are generally observed as positive for health (Valdes et al. 2018), other studies have suggested that overproduction may promote obesity, possibly resulting from greater energy accumulation (Schwiertz et al. 2010; Rahat-Rozenbloom et al. 2014; Teixeira et al. 2013). Indeed, a previous study observed greater fecal SCFA concentrations to be linked with obesity, increased gut permeability, metabolic dysregulation, and hypertension in a human cohort (de la Cuesta-Zuluaga, Mueller, et al. 2018).

The conflicting obesity role of SCFAs identified by existing studies may result from the variation in the gut microbiota, which is shaped by lifestyle and diet. Adequately powered studies in well-characterized populations may permit more rigorous assessments of individual differences. Prior comparative epidemiological studies have broadly focused on either contrasting the gut microbiota of extremely different populations, such as the traditional hunter-gatherers and urban-westernized countries, or ethnically homogenous populations (Pasolli et al. 2019; He et al. 2018; Peters et al. 2018; Zhernakova et al. 2016). Demographic factors represent one of the largest contributors to the individualized nature of the gut microbiome (Falony et al. 2016; Zhernakova et al. 2016; Yatsunenko et al. 2012). Thus, it is unclear to what extent the associations between gut microbiome, SCFA and obesity generalize across different geographies and this, additionally limits our understanding and interpretation, especially when considering the substantial geographic disparities in obesity.

The five diverse, well-defined cohorts from the Modeling the Epidemiologic Transition Study (METS) offers a unique opportunity to examine the issues since they are more representative of most of the world’s population. METS has longitudinally followed an international cohort of approximately 2,500 African-origin adults spanning the epidemiologic transition from Ghana, South Africa, Jamaica, Seychelles, and the US since 2010 to investigate differences in health outcomes utilizing the framework of the epidemiologic transition. Pioneering microbiome studies from the METS cohorts reveal that cardiometabolic risk factors including obesity is significantly associated with reduced microbial diversity, and the enrichment of specific taxa and predicted functional traits in a geographic-specific manner (Dugas, Bernabé, et al. 2018; Fei et al. 2019). While yielding valuable descriptions of the connections between the gut microbiota ecology and disease, particularly obesity, as well as pioneering the efforts of microbiome studies of populations of African-origin on different stages of the ongoing nutritional epidemiologic transitions, these studies, however, have applied small sample size (N=100 to N=655), and also did not utilize all the countries in the METS cohort. Thus, uncertainties remain as to the precise interpretation of the microbiome-obesity associations, which hampers further progress towards diagnostic and clinical applications.

Our new study METS-Microbiome investigated associations between the gut microbiota composition and functional patterns, concentrations of fecal SCFAs and obesity in a large (*N* = 1,934) adult population cohort of African-origin, comprised of Ghana, South Africa, Jamaica, Seychelles, and the US spanning the epidemiologic transition (Dugas, Lie, et al. 2018; Luke et al. 2011). The central hypothesis is that shifts towards the highest end of the epidemiologic transition spectrum is associated with alterations in microbiota diversity and community composition, reductions in levels of fecal SCFAs and obesity.

## Materials and Methods

### Study Cohort

Since 2010, METS, and the currently funded METS-Microbiome study has longitudinally followed an international cohort of African-origin adults spanning the epidemiologic transition from Ghana, South Africa, Jamaica, Seychelles, and US (Dugas, Lie, et al. 2018; Luke et al. 2011). METS utilizes the framework of the epidemiologic transition to investigate differences in health outcomes based on country of origin. The epidemiologic transition is defined using the United Nations Human Development Index (HDI) as an approximation of the epidemiologic transition. Ghana represents a lower-middle income country, South Africa represents a middle-income country, Jamaica and Seychelles represent high income countries and the US represents a very high-income country. This framework has allowed us to investigate aspects of increased Westernization throughout the world (ex. increased consumption of ultra-processed foods) are related to increased prevalence of obesity, diabetes and cardiometabolic diseases. Our data from the original METS cohort demonstrate that the epidemiologic transition has altered habitual diets in the international METS sites, and that reduced fiber intake is associated with higher metabolic risk, inflammation, and obesity across the epidemiologic transition (Mehta et al. 2021). Originally, 2,506 African-origin adults (25–45 yrs), were enrolled in METS between January 2010 and December 2011 and followed on a yearly basis. In 2018, METS participants were recontacted and invited to participate in METS-Microbiome. Participants were excluded from participating in the original METS study if they self-reported an infectious disease, including HIV-positive individuals, pregnancy, breast-feeding or any condition which prevented the individual from participating in normal physical activities. METS-Microbiome was approved by the Institutional Review Board of Loyola University Chicago, IL, US; the Committee on Human Research Publication and Ethics of Kwame Nkrumah University of Science and Technology, Kumasi, Ghana; the Research Ethics Committee of the University of Cape Town, South Africa; the Board for Ethics and Clinical Research of the University of Lausanne, Switzerland; and the Ethics Committee of the University of the West Indies, Kingston, Jamaica. All study procedures were explained to participants in their native languages, and participants provided written informed consent after being given the opportunity to ask any questions.

### Participant anthropometry, sociodemographic and biochemical measurements

Participants completed the research visits at the established METS research clinics located in the respective communities (Luke et al. 2011). Briefly, they presented themselves at the site-specific research clinic early in the morning, following an overnight fast. The weight of the participant was measured without shoes and dressed in light clothing to the nearest 0.1 kg using a standard digital scale (Seca, SC, USA). Height was measured using a stadiometer without shoes and head held in the Frankfort plane to the nearest 0.1 cm. Waist circumference was measured to the nearest 0.1 cm at the umbilicus, while hip circumference was measured to the nearest 0.1 cm at the point of maximum extension of the buttocks. Adiposity (% body fat) was assessed using BIA (Quantum, RJL Systems, Clinton Township, MI), and study specific equations (Luke et al. 2011). Blood pressure was measured using the standard METS protocol using the Omron Automatic Digital Blood Pressure Monitor (model HEM-747Ic, Omron Healthcare, Bannockburn, IL, USA), with the antecubital fossa at heart level. Participants were asked to provide a fecal sample using a standard collection kit (EasySampler stool collection kit, Alpco, NH). Fecal samples were placed within a -80° freezer immediately upon receipt at all the sites. Participants were requested to fast from 8 pm in the evening prior to the clinic examination, during which fasting capillary glucose concentrations were determined using finger stick (Accu-check Aviva, Roche).

### Fecal Short Chain Fatty Acid quantification

As in our previous studies (Nooromid et al. 2020; Lewandowski et al. 2021; Reiman, Layden, and Dai 2021; Barengolts et al. 2019; Navarro et al. 2018; Dugas, Bernabé, et al. 2018), fecal SCFAs were measured using LC-MC/MS at the University of Illinois-Chicago Mass Spectrometry Core using previously published methods (Moreau et al. 2003; Richardson et al. 1989). The LC-MC/MS analysis was completed on an AB Sciex Qtrap 5500 coupled to Agilent UPLC/HPLC system. All samples were analyzed by Agilent poroshell 120 EC-C18 Column, 100Å, 2.7 µm, 2.1 mm X 100 mm coupled to an Agilent UPLC system, which was operated at a flow rate of 400 µl/min. A gradient of buffer A (H_2_0, 0.1% Formic acid) and buffer B (Acetonitrile, 0.1% Formic acid) were applied as: 0 min, 30% of buffer B; increase buffer B to 100% in 4 min; maintain B at 100% for 5 min. The column was then equilibrated for 3 min at 30% B between the injections with the MS detection is in negative mode. The MRM transitions of all targeted compounds include the precursor ions and the signature production ion. Unit resolution is used for both analyzers Q1 and Q3. The MS parameters such as declustering potential, collision energy and collision cell exit potential are optimized in order to achieve the optimal sensitivity. SCFAs are presented as individual SCFAs (μg/g), including: butyric acid, propionic acid, acetic acid and valeric acid, as well as total SCFAs (sum of 4).

METS data showed Ghanaians consumed the greatest amount of both soluble and insoluble fiber and had the lowest percentage energy from fat (42.5% of the Ghanaian cohort, dietary fiber intake: 24.9 g ± 9.7g/day). The US has the highest proportion of energy from fat and the lowest fiber intake of the five sites (3.2% of the US cohort, dietary fiber intake: 14.2 g ± 7.1 g/day).

### DNA extraction, Amplicon Sequencing

Fecal samples were shipped on dry ice to the microbiome core sequencing facility, University of California, San Diego for 16S rRNA gene processing. Fecal samples were randomly sorted, transferred to 96-well extraction plates and DNA was extracted using MagAttract Power Microbiome kit. Blank controls and mock controls (ZymoBiomics) were included per extraction plate, which were carried through all downstream processing steps. Extracted DNA was used for amplification of the V4 region of the 16S rRNA gene with 515F-806R region-specific primers according to the Earth Microbiome Project (Thompson et al. 2017; Walters et al. 2016). Purified amplicon libraries were sequenced on the Illumina NovaSeq platform to produce 150 bp forward and reverse reads through the IGM Genomics Center, University of California San Diego. Full DNA extraction, amplification, quantification, and sequencing protocols and standards are available at http://www.earthmicrobiome.org/protocols-and-standards; (Thompson et al. 2017).

### Bioinformatic analysis

The generated raw sequence data were uploaded and processed in Qiita (Gonzalez et al. 2018) (Qiita ID 13512) an open-source, web-enabled microbiome analysis platform. Sequences were demultiplexed, quality filtered, trimmed, erroneous sequences were removed, and amplicon sequence variants (ASVs) were defined using Deblur (Amir et al. 2017). The deblur ASV table was exported to Qiime2 (Bolyen et al. 2019; Bokulich et al. 2018) and representative sequences of the ASVs were inserted into the Greengenes 13.8 99% identity tree with SATé-enabled phylogenetic placement (SEPP) using q2-fragment-insertion (Bolyen et al. 2019; Mirarab, Nguyen, and Warnow 2012) to generate an insertion tree for diversity computation. Additionally, the deblur ASV table was assigned taxonomic classification using the Qiime2 feature-classifier, with Naive Bayes classifiers trained on the SILVA database (version 138; (McLaren 2020)). A total of 463,258,036 reads, 154,952 ASVs and 1902 samples were obtained from the deblur table. The resulting ASV count table, taxonomy data, insertion tree, and sample metadata were exported and merged into a phyloseq (McMurdie and Holmes 2013) object in R (R Foundation for Statistical Computing, Vienna, Austria) for downstream analysis. Features with less than ten reads in the entire dataset and samples with fewer than 6,000 reads were removed from the phyloseq object. In addition, mitochondrial and chloroplast-derived sequences, non-bacterial sequences, as well as ASVs that were unassigned at phylum level were filtered prior to analyses. There were 433,364,873 reads and 13254 ASVs in the remaining 1873 fecal samples in the phyloseq object. The remaining samples after filtering were rarefied to a depth of 6,000 reads to avoid sequencing bias, before generating alpha diversity measures, leaving 9917 ASVs across 1873 samples.

### Diversity and differential proportional analyses

Alpha diversity measures based on Observed Amplicon Sequence Variants (ASVs), Faith’s Phylogenetic Diversity, and Shannon Index were conducted on rarified samples using phyloseq (McMurdie and Holmes 2013) and picante (Kembel et al. 2010) libraries. Beta diversity was determined using both weighted and unweighted UniFrac distance matrices (Lozupone and Knight 2005), generated in phyloseq. For differential abundance analysis, samples were processed to remove exceptionally rare taxa. First, the non-rarefied reads were filtered to remove samples with < 10,000 reads. Next, ASVs with fewer than 50 reads in total across all samples and/or were present in less than 2% of samples were excluded. This retained 2061 ASVs across 1694 samples. The retained ASVs were binned at genus level, and subsequently used in the analysis of compositions of microbiomes with bias correction (ANCOMBC; (H. Lin and Peddada 2020) to determine specific taxa differentially abundant across sites or obese phenotype. ANCOM-BC is a statistical approach that accounts for sampling fraction, normalizes the read counts by a process identical to log-ratio transformations while controlling for false discovery rates and increasing power. This method applies a library-specific offset term estimated from the observed abundance, which is incorporated into a linear regression model, providing the bias correction. Site, age, sex, BMI were added as covariates in the ANCOM-BC formula to reduce the effect of confounders. The *Bacteroides Prevotella* ratio was calculated by dividing the abundance of the genera *Bacteroides* by *Prevotella*. Participants were classified into *Bacteroides* enterotype (B-type) if the ratio was greater than 1, otherwise *Prevotella* enterotype (P-type).

### Random forest classifier

Random Forest supervised learning models implemented in Qiime2 were used to estimate the predictive power of microbial community profiles for site and obese phenotype. The classifications were done with 500 trees based on 10-fold cross-validation using the QIIME “sample-classifier classify-samples” plugin (Bokulich et al. 2018). A randomly drawn 80% of samples were used for model training, whereas the remaining 20% were used for validation. Further, the 30 most important ASVs for differentiating between site or obese phenotype were predicted and annotated.

### Predicted metabolic gene pathway analysis

The functional potential of microbial communities was inferred using the Phylogenetic Investigation of Communities by Reconstruction of Unobserved States 2 (PICRUSt2) v2.5.1 with the ASV table processed to remove exceptionally rare taxa and the representative sequences as input files (Douglas et al. 2020). The metabolic pathway from the PICRUSt2 pipeline was annotated using the MetaCyc database (Caspi et al. 2016). The predicted MetaCyc abundances (unstratified pathway abundances) were analyzed with ANCOM-BC to determine differentially abundant pathway associations across sites and obese status. Site, age, sex, BMI were added as covariates in the ANCOM-BC formula to reduce the effect of confounders.

### Statistical Analysis

All statistical analyses and graphs were done with R software. Kruskal-Wallis test and Permutational Analysis of Variance (PERMANOVA) test with 999 permutations using the Adonis function in the vegan package (Oksanen et al. 2013) were performed to compare alpha and beta diversity measures respectively with multiple groups comparison correction. PERMANOVA models were adjusted for BMI, age, sex for country whereas age, sex and country were accounted for in obese groups. Variables that showed significant differences in the PERMANOVA analyses, PERMDISP test was performed to assess differences in dispersion or centroids. For differential abundance analysis, the false-discovery rate (FDR) method incorporated in the ANCOM-BC library was used to correct P values for multiple testing. A cut-off of P_adj_ < 0.05 was used to assess significance. Spearman correlations were performed between concentrations of short chain fatty acids, Shannon diversity or concentrations of short chain fatty acids and differentially abundant taxa that were identified either among study sites or in obese and non-obese individuals. The resulting p-values were adjusted for multiple testing using the false-discovery rate (FDR). P value < 0.05 was considered statistically significant. A mixed model was built using lme4 package to assess whether total SCFAs could be predicted by Shannon diversity, obesity, and country, setting obesity and Shannon diversity as fixed effects and random intercept by country.

### Data availability

All 16S rRNA gene sequence data are publicly available via the QIITA platform (https://qiita.ucsd.edu) under the study identifier (ID=13512) and will soon be available on the European Bioinformatics Institute (EBI) site.

## Results

### Obesity differs significantly across the epidemiological transition

From 2018-2019, the METS-Microbiome study recruited 2,085 participants (∼60% women) ages 35-55 years old from five different sites (Ghana, South Africa, Jamaica, Seychelles, and US). Of these participants, 1,249 have been followed on a yearly basis since 2010 under the parent METS study. Data from 1,867 participants with complete data sets were used in this analysis. Overall mean age was 42.5 ± 8.0 years (**Table 1**). Mean fasted blood glucose was 105.2 ± 39.4 mg/dL, mean systolic blood pressure was 123.4±18.1 mm Hg and mean diastolic blood pressure was 77.2 ± 13.1 (**Table 1**). When compared to the high-income countries (Jamaica, Seychelles, and US), both women and men from the lower- and middle-income countries (Ghana and South Africa) had significantly lower BMI, fasted blood glucose and blood pressure (systolic and diastolic). Mean BMI was lowest in the South African men (22.3 kg/m2 ± 4.1) and highest in US women (36.3 kg/m2 ± 8.8). When compared to the US, all sites had significantly lower prevalence of obesity (p<0.001 for all sites except for Seychelles: p=0.02). Prevalence of hypertension was lowest in Ghanaian men (33.1%) and highest in US men (72.7%). Prevalence of diabetes was lowest in South African women and men (3.5% for women and men) and highest for Seychellois men (22.8%). When compared to the US, prevalence of hypertension and diabetes was significantly lower in countries at the lower end of the spectrum of HDI (i.e., Ghana and South Africa) when compared to the US (p<0.001).

**Table 1.** METS-Microbiome participant characteristics from Ghana, South Africa, Jamaica, Seychelles and US

| **Women** |  |  |  |  |  |
| --- | --- | --- | --- | --- | --- |
|  | **Ghana** | **South Africa** | **Jamaica** | **Seychelles** | **US** |
|  | n=254 | n=228 | n=263 | n=196 | n=213 |
| *Age (years)* | 40.74 ± 8.1 | 35.56 ± 7.8 | 45.16 ± 7.5 | 43.84 ± 6.1 | 45.44 ± 6.4 |
| *BMI (kg/m<sup>2</sup>)* | 28.30 ± 5.9 | 33.42 ± 8.6 | 32.12 ± 7.3 | 30.32 ± 7.2 | 36.34 ± 8.8 |
| *Obese (%)* | 45,0% | 61,0% | 60,4% | 49,5% | 74,7% |
| *SBP (mm Hg)* | 117.1 ± 18.5 | 115.20 ± 17.1 | 126.08 ± 19.0 | 123.28 ± 17.8 | 124.19 ± 18.4 |
| *DBP (mm Hg)* | 70.53 ± 12.2 | 75.20 ± 12.1 | 79.41 ± 12.6 | 79.37 ± 14.4 | 81.52 ± 12.1 |
| *Hypertensive (%)* | 37,5% | 37,3% | 57,4% | 55,5% | 65,4% |
| *Glucose (mg/dL)* | 110.45 ± 62.7 | 89.17 ± 20.0 | 107.46 ± 39.1 | 111.35 ± 27.2 | 107.07 ± 44.0 |
| *Diabetic (%)* | 10,0% | 3,5% | 12,9% | 13,9% | 19,9% |
| **Men** |  |  |  |  |  |
|  | **Ghana** | **South Africa** | **Jamaica** | **Seychelles** | **US** |
|  | n=117 | n=171 | n=133 | n=164 | n=107 |
| *Age (years)* | 43.92 ± 8.7 | 36.53 ± 7.2 | 44.42 ± 7.5 | 44.57 ± 5.1 | 47.12 ± 5.5 |
| *BMI (kg/m<sup>2</sup>)* | 23.7 ± 4.4 | 22.26 ± 4.1 | 24.8 ± 5.3 | 28.46 ± 5.5 | 30.37 ± 8.2 |
| *Obese (%)* | 13,4% | 5,3% | 15,7% | 39,2% | 44,4% |
| *SBP (mm Hg)* | 121.28 ± 15.4 | 122.71 ± 15.5 | 129.23 ± 17.1 | 130.43 ± 16.2 | 130.67 ± 16.0 |
| *DBP (mm Hg)* | 68.02 ± 13.0 | 75.32 ± 11.1 | 78.07 ± 11.5 | 81.64 ± 12.1 | 82.37 ± 12.2 |
| *Hypertensive (%)* | 33,1% | 45,0% | 50,3% | 65,9% | 72,7% |
| *Glucose (mg/dL)* | 100.52 ± 19.4 | 94 ± 23.4 | 99.04 ± 33.1 | 124.26 ± 44.2 | 107 ± 36.2 |
| *Diabetic (%)* | 4,6% | 3,5% | 4,8% | 22,8% | 17,5% |

### Microbial community composition and predicted metabolic potential differs significantly between countries and correlates with obesity

Following the removal of control samples and those that had fewer than 6,000 reads and features less than ten reads in the entire dataset, a total of 433,364,873 16S rRNA gene sequences were generated from the 1,873 fecal samples which were clustered into 13,254 ASVs. Country of origin describes most of the variation in microbial diversity and composition, with significant differences in both alpha and beta diversity. Although there were major variations in alpha diversity between countries and large degree of inter-individual variation within countries, Ghana showed significantly greater diversity for all the alpha diversity metrics (Observed ASVs, Shannon Diversity and Faith’s phylogenetic diversity) when compared to all other countries. The Seychelles and US had the lowest alpha diversity (**Figure 1**, **Table 2**). The stool microbiota alpha diversity of non-obese individuals was significantly greater when compared with that of obese individuals (**Figure 1**). Beta diversity was also significantly different between countries (**Figure 1**, **Table 3 & Supplementary Table 2;** principal coordinate analysis, weighted UniFrac distance; F-statistic =58.67; p < 0.001; unweighted UniFrac distance; F= 39.87; p < 0.001) and obese group (weighted UniFrac distance; F-statistic =2.39; p = 0.031; unweighted UniFrac distance; F=6.06; p < 0.001).

**Figure 1.**
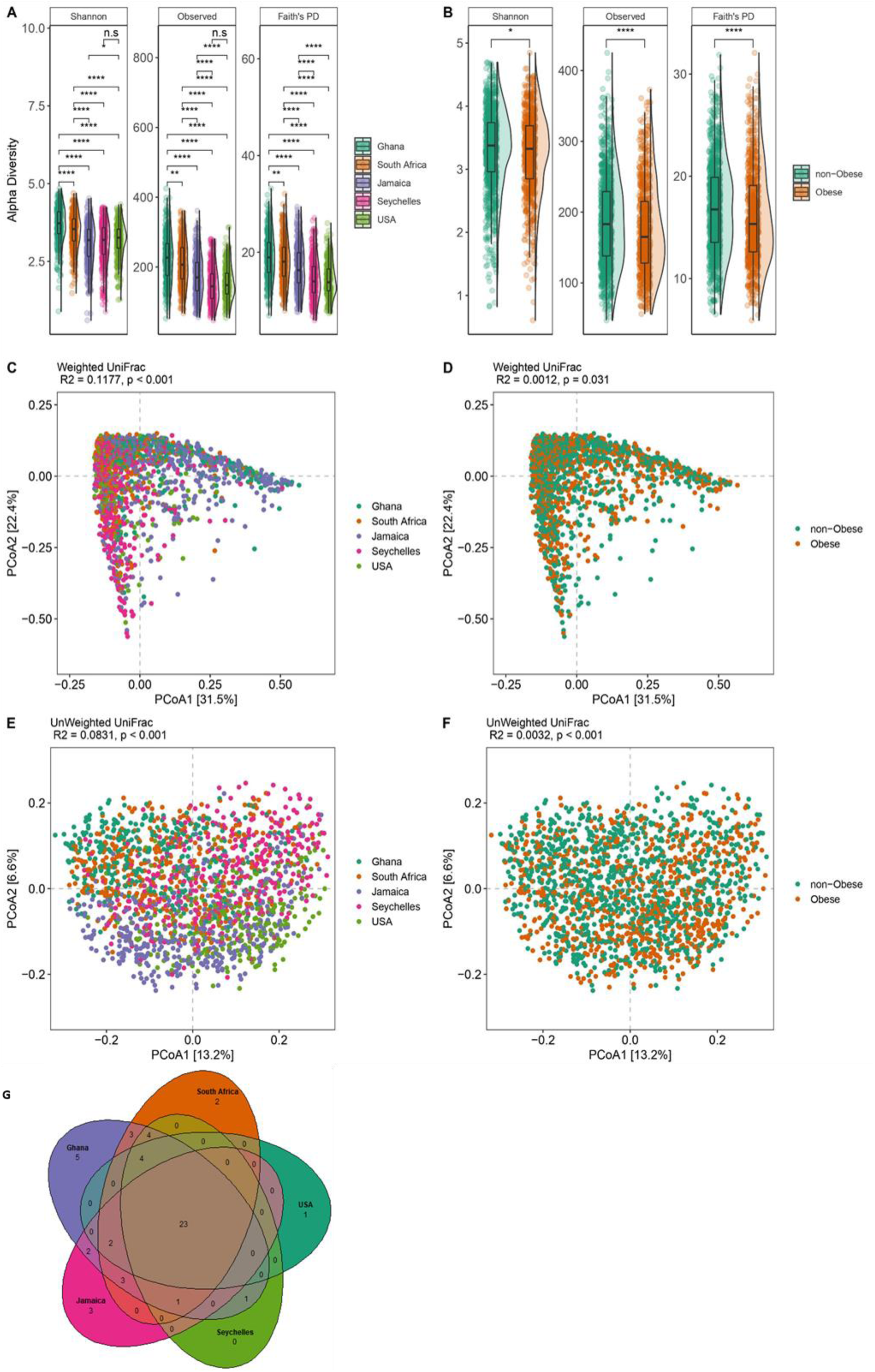
Variation in gut microbiome diversity and composition. (**A)** Alpha diversity estimated by Shannon, Observed ASVs and Faith’s PD (Phylogenetic Diversity) between countries. (**B**) Alpha diversity estimated by Shannon, Observed ASVs and Faith’s PD (Phylogenetic Diversity) between obese and non-obese. *p < 0.05, ****p < 0.0001 Alpha diversity metrics (Faith’s PD, Observed ASVs and Shannon) are shown on the y-axis in different panels, while country or obese group are shown on the x-axis. (**C**) Beta diversity principal coordinate analysis based on weighted UniFrac distance between countries. (**D**) Beta diversity principal coordinate analysis based on weighted UniFrac distance between obese and non-obese. (**E**) Beta diversity principal coordinate analysis based on unweighted UniFrac distance between countries. (**F**) Beta diversity principal coordinate analysis based on unweighted UniFrac distance between obese and non-obese. Proportion of variance explained by each principal coordinate axis is denoted in the corresponding axis label. (**G**) Venn diagram of shared and unique ASVs between the five countries. Statistical significance adjusted for multiple comparisons using false discovery rate (FDR) correction are indicated: *, P < 0.05; **, P < 0.01; ***, P < 0.001; ***, P < 0.001 across countries and obese groups (Kruskal-Wallis test) for alpha diversity or by permutational multivariate analysis of variance (PERMANOVA) for beta diversity.

**Table 2.** Alpha diversity estimated by Shannon, Observed ASVs and Faith’s PD (Phylogenetic Diversity) between countries and obesity status. q-value are FDR-corrected p values representing statistical significance (p<0.05) of alpha diversity metrics between the countries. Data are presented by median (interquartile range). FDR = False Discovery Rate

|  |  | **N** | **Faith's PD** | **Shannon** | **Observed** |
| --- | --- | --- | --- | --- | --- |
| **Ghana** | Obese | 243 | 19.2(16.2-21.8) | 3.73(3.41,4.09) | 228(184,267) |
|  | Non-Obese | 89 | 17.9(14.6,21.6) | 3.69(3.30,4.05) | 217(155,252) |
| **South Africa** | Obese | 208 | 17.2(14.0,19.9) | 3.21(2.69,3.52) | 174(138,212) |
|  | Non-Obese | 179 | 15.9(13.1,19.9) | 3.12(2.65,3.54) | 165(126,216) |
| **Jamaica** | Obese | 217 | 14.2(11.6,17.2) | 3.2(2.72,3.56) | 146(110,184) |
|  | Non-Obese | 147 | 13.4(11.5,16.5) | 3.15(2.69,3.56) | 136(108,173) |
| **Seychelles** | Obese | 233 | 18(15.1,20.4) | 3.51(3.14,3.79) | 204(166,246) |
|  | Non-Obese | 141 | 18.6(15.0,22.4) | 3.62(3.21,3.98) | 212(165,269) |
| **US** | Obese | 112 | 13.6(12.1,16.6) | 3.23(2.86,3.50) | 144(125,180) |
|  | Non-Obese | 195 | 13.9(12.1,16.5) | 3.3(3.00,3.57) | 150(122,184) |
| **p-value** |  |  | <0.001 | <0.001 | <0.001 |
| **q-value** |  |  | <0.001 | <0.001 | <0.001 |

Median (IQR)
p-value: Kruskal-Wallis rank sum test
q-value: False discovery rate correction for multiple testing

**Table 3.** Adjusted Multivariate Analysis for the entire cohort and by each country. Statistical significance from permutational multivariate analysis of variance (PERMANOVA) test, *p* < 0.05. All *p*-values are generated based on 999 permutations

|  | Overall |  | Ghana |  | South Africa |  | Jamaica |  | Seychelles |  | US |  |
| --- | --- | --- | --- | --- | --- | --- | --- | --- | --- | --- | --- | --- |
|  | R<sup>2</sup> | P | R<sup>2</sup> | P | R<sup>2</sup> | P | R<sup>2</sup> | P | R<sup>2</sup> | P | R<sup>2</sup> | P |
| **Obese** | 0.003 | 0.001 | 0.004 | 0.032 | 0.007 | 0.002 | 0.002 | 0.732 | 0.003 | 0.279 | 0.004 | 0.154 |
| **Sex** | 0.003 | 0.001 | 0.005 | 0.018 | 0.007 | 0.002 | 0.009 | 0.009 | 0.01 | 0.001 | 0.01 | 0.001 |
| **Age** | 0.001 | 0.135 | 0.113 | 0.471 | 0.083 | 0.708 | 0.102 | 0.062 | 0.063 | 0.576 | 0.094 | 0.252 |
| **Country** | 0.083 | 0.001 |  |  |  |  |  |  |  |  |  |  |

Next, we compared fecal microbiota diversity between obese individuals with their non-obese counterparts within each country independently (**Supplementary Table 1**). Greater alpha diversity was detected in non-obese subjects in the Ghanaian (Observed ASVs, Faith PD; p<0.05) and South African cohorts (Observed ASVs; p<0.05) only. Similarly, significant differences in beta diversity between obese and non-obese microbiota were observed in Ghana (Unweighted UniFrac; p<0.05), South Africa (Unweighted UniFrac; p<0.05) and US (Weighted UniFrac; p<0.05) data sets (**Table 3 & Supplementary Table 2**). These results suggest that the beta diversity differences observed in the Ghanaian and South African participants may partly be due to the presence of more abundant fecal microbiota taxa in the fecal samples whereas among the US participants, the differences may be related to the abundance of rare taxa. Collectively, these observations suggest that country is a major driver of the variance in gut microbiota diversity and composition among participants with or without obesity with marked contributions from Ghana and South Africa and modest contribution from the US in the overall cohort.

We also examined whether country of origin or obesity relates to the presence of specific microbial genera frequently used to stratify humans into enterotypes (Arumugam et al. 2011). As expected, large differences in enterotype between the countries were observed. The *Prevotella* enterotype (P-type) was enriched on the African continent, with 81% and 62% in Ghanaians and South Africans respectively while *Bacteroides* enterotype (B-type) was dominant in the US (75%), Jamaican cohorts (68%), and comparable proportions of both enterotypes among individuals from Seychelles. Further, obese individuals displayed a greater abundance of B-type whereas a higher proportion of the P-type associated with the non-obese group (**Supplementary Table 3**). Consistent with this observation, the abundance of B-type correlated with higher BMI (p=0.004) than P-type. Significantly greater diversity and increased levels of total SCFA were observed in participants in the P-type (**Supplementary Table 3**). The relative abundance of shared and unique features between the different countries illustrated by the Venn diagram showed that Ghana carries the largest proportion of unique taxa than the other countries, and US the lowest (**Figure 1**).

### Microbial taxa differ significantly between countries and between lean and obese individuals

In comparison with the US, South African fecal microbiota had a significantly greater proportion of *Clostridium*, *Olsenella*, Bacilli and *Mogibacterium*; Jamaican samples had a significantly greater proportion of Bacilli, *Bacteroides*, Clostridia, *Dialister*, Enterobacteriaceae, and Oscillospiraceae; Seychelles samples had a significantly greater proportion of *Clostridium*, *Olsenella* and *Haemophilus*; and Ghanaian samples had a significantly greater proportion of *Clostridium*, *Prevotella*, *Weisella,* Enterobacteriaceae and Butyricicoccaceae. The US samples had a significantly greater proportion of *Aldercreutzia*, *Anaerostipes*, *Clostridium*, *Eggerthella*, *Eisenbergiella*, Ruminococcaceae and *Sellimonas* compared to the 4 countries (**Supplementary Figure 1**).

When adjusted for country, age, and sex (p < 0.05; false discovery rate (fdr)-corrected), 38 Amplicon Sequence Variants (ASVs) were significantly different between obese and non-obese groups. The obese group was characterized by an increased proportion of *Allisonella*, *Dialister*, *Oribacterium*, *Mitsuokella*, and *Lachnospira*, whereas non-obese microbiota had a significantly greater proportion of *Alistipes*, *Bacteroides*, *Clostridium*, *Parabacteroides*, *Christensenella*, *Oscillospira*, Ruminococcaceae (UBA1819), and Oscillospiraceae (UCG010) (**Supplementary Figure 1**).

### Microbial taxonomic features predict obesity overall and within each country

Using supervised Random Forest machine learning, the predictive capacity of the gut microbiota features in stratifying individuals to country of origin, sex, or with metabolic phenotypes were assessed. The predictive performance of the model was calculated by area under the receiver operating characteristic curve (AUC) analysis, which showed a high accuracy for country of origin (AUC = 0.97), and a comparatively lower level of predictive accuracy for obese state (AUC = 0.65) (**Figure 2**). Sex was predicted with AUC = 0.75, the diabetes status with AUC = 0.63, hypertensive status with AUC = 0.65 and glucose status with AUC = 0.66. Random Forest analysis was also used to identify the top 30 microbial taxonomic features that differentiate between countries and obese states. Similar to the ANCOMBC results, *Prevotella* and *Streptococcus* were at a greater proportion in the microbiota of Ghanaian and non-obese individuals, whereas *Mogibacterium* was at a greater proportion in the South African cohort. A greater proportion of *Megasphaera* was associated with the Jamaican cohort, while a greater proportion of Ruminococcaceae was observed in the American microbiota. *Weisella*, which was identified as having a significantly greater proportion in the Ghanaian cohort using ANCOMBC, was observed to be a discriminatory feature for Seychelles microbiota using Random Forest (**Supplementary Figure 2**).

**Figure 2.**
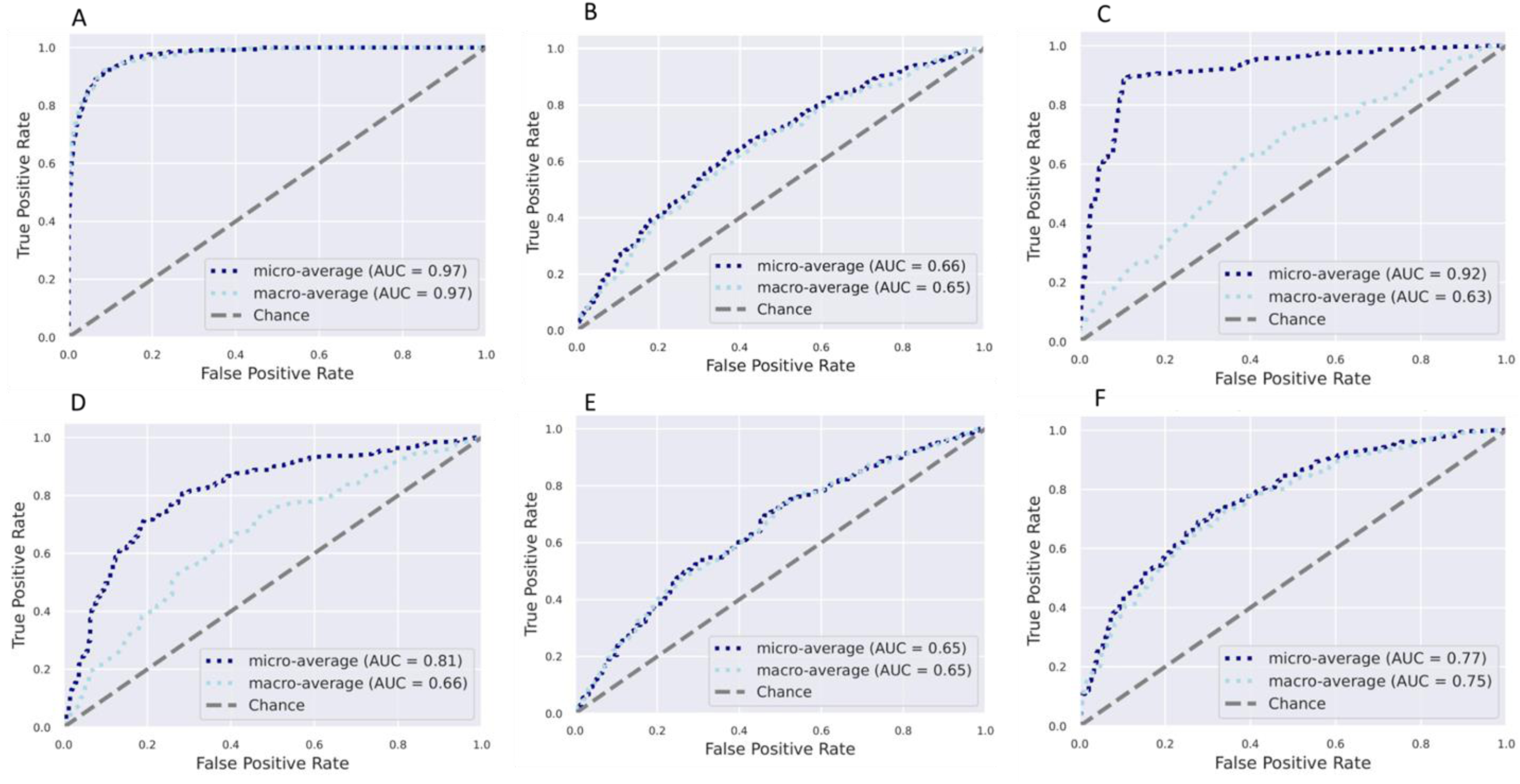
Receiver operating characteristic curves showing the classification accuracy of gut microbiota in a Random Forest model. Classification accuracy for estimating (**A**). All countries; (**B**) Obesity status, (**C**). Diabetes status; (**D**). Glucose status; (**E**). Hypertensive status; (**F**). Sex are presented. AUC= area under the curve

Similarly, the predictive capacity of the gut microbiota features in stratifying individuals by obese state was assessed at each of the five study sites. The predictive performance of the model was calculated by AUC analysis, which showed a moderate accuracy for obese state for all sites, namely, Ghana (AUC = 0.57), South Africa (AUC = 0.52), Jamaica (AUC = 0.48), Seychelles (AUC = 0.43) and US (AUC = 0.52) (**Supplementary Figure 3**).

### Predicted genetic metabolic potential differs by country and obesity status

The predicted potential microbial functional traits resulting from the compositional differences in microbial taxa between countries and obese state were assessed. PICRUSt2 predicted a total of 372 MetaCyc functional pathways. ANCOM-BC analysis adjusted for sex, age and BMI identified 67 pathways (p< 0.05; false discovery rate (fdr)-corrected), LFC>1.4) that accounted for discriminative features between the 4 different countries with the US (**Supplementary Figure 4**). In comparison with US, MetaCyc pathways differentially increased in Ghana and Jamaica include methylgallate degradation, norspermidine biosynthesis (PWY-6562), gallate degradation I pathway, gallate degradation II pathway, histamine degradation (PWY-6185), and toluene degradation III (via p-cresol) (PWY-5181). South African samples had a greater proportion of L-glutamate degradation VIII (to propanoate) (PWY-5088), isopropanol biosynthesis (PWY-6876), creatinine degradation (PWY-4722), adenosyl cobalamin biosynthesis (anaerobic) (PWY-5507), respiration I (cytochrome c) (PWY-3781). MetaCyc pathways linked to norspermidine biosynthesis (PWy-6562), mycothiol biosynthesis (PWY1G-0), were at a greater proportion in the Seychelles samples, whereas reductive acetyl coenzyme A (CODH-PWY), and chorismate biosynthesis II (PWy-6165) were depleted in the US samples. ANCOM-BC analysis adjusted for site, sex and age identified 24 predicted pathways that differentiated between obese and non-obese individuals (**Supplementary Figure 4**). Notably, the microbiota of non-obese individuals had a greater proportion of predicted pathways including the TCA cycle, amino acid metabolism (P162-PWY, PWY-5154, PWY-5345), ubiquinol biosynthesis-related pathways (PWY-5855, PWY-5856, PWY-5857, PWY-6708, UBISYN-PWY), cell structure biosynthesis and nucleic acid processing (PWY0 845, PYRIDOXSYN-PWY).

Next, KEGG orthology (KO) involved in pathways related to butanoate (butyrate) metabolism and LPS biosynthesis were investigated. Predicted genes involved in butyrate biosynthesis pathways showed that enoyl-CoA hydratase enzymes (K01825, K01782, K01692), lysine, glutarate /succinate enzymes (K07250, K00135, K00247), glutarate/Acetyl CoA enzymes (K00175, K00174, K00242, K00241 K01040, K01039) were differentially abundant in participants from Ghana, South Africa, Jamaica, and Seychelles in comparison to the US cohort. The relative abundance of succinic semialdehyde reductase (K18122) was significantly increased only in South Africa, Jamaica, and Seychelles population. Further, predicted genes proportionally abundant only in specific countries were observed. For instance, succinate semialdehyde dehydrogenase (K18119) was enriched only in the Ghanaian cohort, 4-hydroxybutyrate CoA-transferase (K18122) enriched among South African participants and lysine/glutarate/succinate enzyme (K14268) differentially abundant within the Seychelles population. The relative abundance of predicted genes encoded for enzymes such as maleate isomerase (K10799), 3-oxoacid CoA-transferase(K01027) and pyruvate/acetyl CoA (K00171, K00172, K00169) were greater in the US participants compared with participants from the 4 countries (**Supplementary Figure 5**). The non-obese exhibited a significantly greater abundance of genes that catalyze the production of butyrate via the fermentation of pyruvate or branched amino-acids such as enoyl-CoA hydratase enzyme (K0182), Leucine/Acetyl CoA enzyme (K01640) and pyruvate/acetyl CoA enzyme (K00171, K00172, K00169, K1907) by contrast obese individuals were differentially enriched for succinyl-CoA:acetate CoA-transferase (K18118) (**Supplementary Figure 5**). All analyses were adjusted for country, sex, BMI and age (fdr-corrected p < 0.05).

Several gut microbial predicted genes involved in LPS biosynthesis differentially enriched among the countries (p< 0.05; false discovery rate (fdr)-corrected) were identified. In particular, the relative abundance of specific LPS genes (K02560, K12973, K02849, K12979, K12975, K12974) were significantly enriched in Ghana, South Africa, Jamaica, and Seychelles when compared with US. Higher proportions of LPS genes including K12981, K12976 K09953, K03280 were significantly increased in Seychelles samples in comparison with US samples and also significantly increased in the US cohorts in comparison with participants from Ghana, South Africa, and Seychelles. US samples had a greater proportion of the following genes (K15669, K09778, K07264, K03273, K03271) in comparison with the other 4 countries (**Supplementary Figure 6**). Non-obese individuals had a greater abundance of predicted genes encoding LPS biosynthesis (K02841, K02843, K03271, K03273, K19353, K02850) whereas only 1 LPS gene (K02841) differentially elevated in the non-obese group (**Supplementary Figure 6)**. All analyses were adjusted for country, sex, BMI and age (fdr-corrected p < 0.05).

### Microbial community composition and predicted metabolic potential correlates with observed fecal SCFA concentrations

All countries had significantly higher weight-adjusted fecal total SCFA levels when compared to the US participants (p<0.001), with Ghanaians having the highest weight-adjusted fecal total SCFA levels (**Supplementary Table 4**). When compared to their obese counterparts, non-obese participants had significantly higher weight-adjusted fecal total and individual SCFA levels (**Supplementary Table 5**). Total SCFA levels displayed weak, but significantly positive correlation with Shannon diversity (r = 0.0.074). A similar trend was observed in the different individual SCFAs, namely valerate (r = 0.19), butyrate (r = 0.12), propionate (r = 0.073) and acetate (r = 0.058) (**Figure 3**). Observed ASVs were not significantly correlated with total SCFAs (p>0.05). Levels of acetate, butyrate and propionate exhibited strong significant correlations with total SCFA, whereas valerate levels significantly correlated negatively (r = -0.09) with total SCFAs. Next, we assessed if levels of total SCFAs could be predicted by a mixed model. Country explained 45.7% of the variation in SCFAs. No significant effect was explained either by obesity or Shannon diversity.

**Figure 3.**
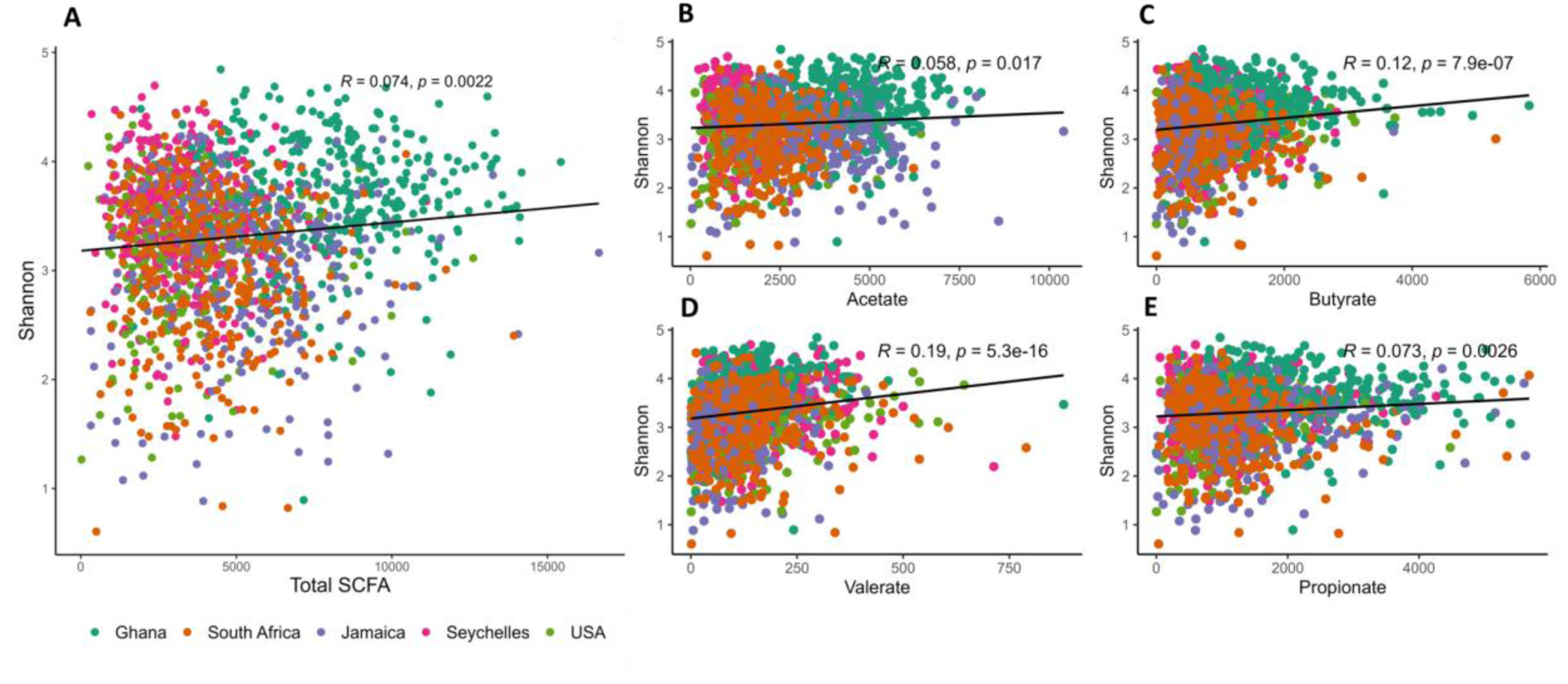
Correlations between alpha diversity and concentrations of the different types of fecal short chain fatty acids (SCFAs) among countries. Shannon index correlates positively with (**A**) total SCFA; (**B**) Acetate; (**C**) Butyrate; (**D**) Propionate; (**E**) Valerate.

### Microbial taxonomy correlates with SCFA concentration and obesity status

To explore the connection between SCFAs with gut microbiota, Spearman correlations between taxa that were proportionally significantly different between countries and concentrations of SCFAs were determined. Valerate negatively correlated with the proportion of *Clostridium*, *Prevotella*, *Faecalibacterium*, *Roseburia* and *Streptococcus*, which were all positively correlated with acetate, propionate, and butyrate. Similarly, the proportions of Christensenellaceae, *Eubacterium*, and UCG 002 (Ruminococcaceae) were significantly positively associated with valerate, and negatively correlated with acetate, propionate, and butyrate. In addition, only a single ASV annotated to *Ruminococcus* was observed to be positively associated with all 4 SCFAs (**Figure 4**). Similarly, Spearman’s rank correlation coefficients were calculated between the differentially abundant ASVs identified between obese and non-obese group with concentrations of SCFAs. Broadly, the proportions of most ASVs were significantly positively associated with acetate in comparison with the other 3 SCFAs. Consistent with the correlations mentioned above, valerate negatively correlated with most ASVs that were found to be positively correlated with the three major SCFAs, acetate, propionate, and butyrate and vice versa. The relative proportions of ASVs belonging to *Allisonella*, Erysipelotrichaceae and *Libanicoccus* positively correlated with acetate, propionate, and butyrate, whereas significantly negative relationships were observed between *Parabacteroides* and *Bacteroides* abundances with the aforementioned SCFAs. Valerate showed significantly positive associations with Oscillospiralles and Ruminococcaceae abundances and significantly negative correlations with *Lachnospira* and *Eggerthella* abundances (**Figure 4**).

**Figure 4.**
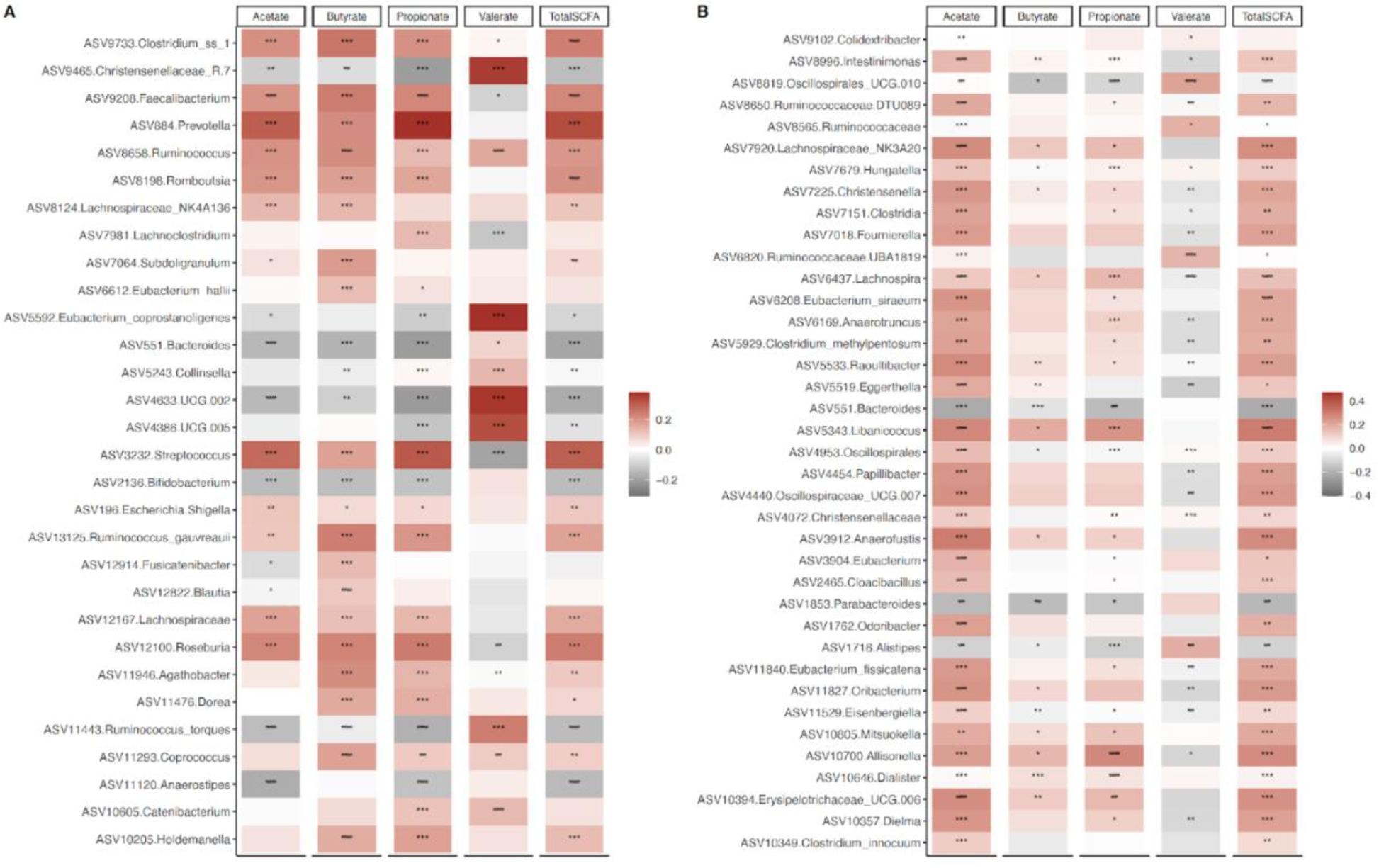
Associations of gut microbiota ASVs with concentrations of short chain fatty acids (SCFAs). (**A**) Heatmap of Spearman’s correlation between concentrations of SCFAs and top 30 differentially abundant ASVs (identified by ANCOM-BC) among countries. (**B**) Heatmap of Spearman’s correlation between concentrations of SCFAs and differentially abundant ASVs (identified by ANCOM-BC) for obese. Correlations are identified by Spearman’s rank correlation coefficient. Brick red squares indicate positive correlation, gray squares represent negative correlation and white squares are insignificant correlation. Mapping from FDR adjusted p values are denoted as: *, ** and ***, corresponding to *p* < 0.05, <0.01 and <0.001 respectively.

## Discussion

By leveraging a well characterized large population-based cohort of African origin residing in geographically distinct regions of Ghana, South Africa, Jamaica, Seychelles, and the US, we examined the relationships between gut microbiota, SCFAs and adiposity. Our data revealed profound variations in gut microbiota, which are reflected in the significant changes in community composition, structure, and predicted functional pathways as a function of population obesity and geography, despite their shared ancestral background. Our data further revealed an inverse relation between fecal SCFA concentrations, microbial diversity, and obesity; importantly, the utility of the microbiota in predicting whether an individual was lean or obese was inversely correlated with the income-level of the country of origin. Overall, our findings are important for understanding the complex relationships between the gut microbiota, population lifestyle and the development of obesity, which may set the stage for defining the mechanisms through which the microbiome may shape health outcomes in populations of African-origin.

It has previously been reported that geographic origin can modulate the composition of the gut microbiota (Yatsunenko et al. 2012; De Filippo et al. 2010, 2017). Accordingly, taxonomic profiling revealed significant differences in gut microbiota richness and diversity among the different countries in a continuum manner. Notably, we detected greater microbiota diversity in Ghana, while depleted microbiota diversity was associated with the US, representing the lowest and the highest end along the epidemiologic transition spectrum respectively, while South Africans, Jamaicans and Seychellois ranked in between. Our findings are consistent with our previous METS studies (Fei et al. 2019; Dugas, Bernabé, et al. 2018) and other large scale continental cohort studies (De Filippo et al. 2010, 2017; Yatsunenko et al. 2012; Schnorr et al. 2014; Clemente et al. 2015; Rampelli et al. 2015; Gomez et al. 2016; Mancabelli et al. 2017), that report a higher bacterial diversity and composition/microbial richness in traditionally non-western groups that distinguish them from urban-industrialized individuals whose diets are low in fiber and high in saturated fats (E. D. Sonnenburg and Sonnenburg 2019; Kolodziejczyk, Zheng, and Elinav 2019). Although we observe enrichment in the relative abundance of several taxa associated with country of origin in our cohorts, we also detect a pattern where the gut microbiota of Ghanaian and South African cohort tends to share many features, while the gut microbiota of the Jamaican cohort shared many features with all 4 countries, possibly reflecting the ongoing epidemiological transitional nature of their communities represented by the overlap with western and traditionally non-western populations. Notably, traditionally non-western associated taxa including *Prevotella*, *Butyrivibrio*, *Weisella* and *Romboutsia* were enriched in participants from Ghana and South Africa, as suggested previously (Mancabelli et al. 2017). Western-associated taxa such as *Bacteroides* and *Parabacteroides* were enriched in individuals from Jamaica and the US (Mancabelli et al. 2017; Kao et al. 2015), while an ASV annotated as *Olsenella* was proportionally abundant in Seychelles microbiota. *Bifidobacterium* and *Aldercreutzia* were enriched in the US cohort. *Clostridium* sensu stricto 1 was over-represented in all 4 countries in comparison with the US. We also found greater enrichment of VANISH taxa including *Butyricicoccus and Succinivibrio* in the Ghanaian cohort, in line with individuals practicing traditional lifestyles (Pasolli et al. 2019). *Prevotella* is usually associated with plant-based diets rich in dietary fibers, while *Bacteroides* abundance broadly correlates with diets high in fat, animal protein, and sugars (Gupta, Paul, and Dutta 2017; Wu et al. 2011), which is in agreement with our enterotype analysis where a Prevotella-rich microbiota dominates the Ghanaian and South African gut, while a Bacteroides-rich microbiota dominated in the high-income countries. *Prevotella* is known to produce high amounts of SCFAs (T. Chen et al. 2017), so its depletion may be associated with the observed concomitant reduction in SCFA concentrations. Increased SCFA synthesis is associated with a reduction in obesity, which is supported by our observations, whereby elevated concentrations of total SCFA and a concomitant reduction in obesity is associated with the *Prevotella* dominated gut of the Ghanaian cohort. Our results support a potential role for geography in reinforcing variations in the gut microbiota in our study cohort despite shared origin. Geography may reflect subtle shifts in lifestyle and/or environmental exposures including heterogeneity of dietary sources, exposure to medications, socioeconomic factors, medical history, and biogeographical patterns in microbial dispersion (Asnicar et al. 2021; Pasolli et al. 2019; Costello et al. 2012; He et al. 2018).

We also inferred the metabolic capacity of the gut microbiota associated with the different countries. Several metabolic pathways linked to carrier, cofactor and vitamin biosynthesis, biosynthesis/degradation of amines, amino acids, aromatic xenobiotics, and tricarboxylic acid (TCA) cycle were differentially enriched between the different countries compared with the US. These pathways are involved in biochemical reactions that regulate several processes including energy metabolism, inflammation, epigenetic processes, and oxidative stress. Several of these observed pathways have been reported in different populations (Yu et al. 2021; Karlsson et al. 2013; N. Qin et al. 2014) indicating that the gut microbiota can directly influence host metabolism, although a majority of these molecules can also be synthesized by the host or supplied through diet. In our cohort, functional shifts observed in participants from Ghana and Jamaica included the enrichment of the metabolic pathway for degradation of gallate. Metabolites generated from the gallate pathway include phenolic catechin metabolites which are thought to alleviate obesity-related pathologies and also promote a healthy and beneficial human gut microbiota composition (Marchesi et al. 2016; Liu et al. 2021). We found pathways related to glutamate degradation which can be fermented to butyrate and propionate enhanced among South Africans and Ghanaians in comparison with the US. In Seychelles, a pathway involved in mycothiol biosynthesis was upregulated. Mycothiol is a protective antioxidant produced by the members of the Actinobacteria phylum and is involved in the removal of toxic compounds from cells (Newton, Buchmeier, and Fahey 2008). The predicted abundance of mycothiol biosynthesis pathway was identified as underrepresented in the microbiome of individuals with depressive symptoms in a South Korean population (S.-Y. Kim et al. 2022).

We further identified increased abundances in pathways related to the generation of SCFAs such as acetyl coenzyme A pathway, threonine biosynthesis and leucine degradation pathway in the microbiomes of all 4 countries in comparison with the US. Threonine can be metabolized to SCFAs acetate and propionate (Davila et al. 2013) and indeed, genes linked with threonine metabolism have been identified in the human gut microbiome (Abubucker et al. 2012). Taken together, our results suggest that the observed country-specific microbial differences and abundances accompany variance in the distribution of functional pathways abundances, although we are unable to ascertain what the sources are that may explain these differences in the predicted functional enrichment due the inherent limitation in functional resolution of 16S rRNA sequence data in PICRUSt2 analysis. Further studies are required to evaluate the potential causal relations of these gut microbial functions with health outcomes using shotgun metagenomic sequencing which offers robust inferences of functional pathways.

Preclinical germ-free mouse models provide early causal links between gut microbial ecology and obesity (Ley et al. 2005; Bäckhed et al. 2007). Thereafter, follow up studies in human cohorts have sought to identify a consistent microbiota signature across populations that can be used to predict obesity. However, identifying obesity-specific microbiome features have proven difficult because the results are often not in agreement (Finucane et al. 2014). Therefore, we sought to examine the fecal levels of individual SCFA types and linking to variations in gut microbiota in obese and non-obese individuals in our large African cohort. The bulk of evidence from prior studies show that obesity is associated with a less diverse bacterial community (Turnbaugh et al. 2009; Dugas, Bernabé, et al. 2018; Peters et al. 2018). Accordingly, we observed that our obese group harbor a significantly lower microbiota diversity and differences in community composition.

Although the mechanism by which the gut microbiota influences obesity are not fully understood, several mechanisms have been proposed. For instance, the regulation of host energy metabolism and body mass concept demonstrate that a perturbed gut bacteria community contributes to the development of obesity by providing excess energy to the host via the fermentation of indigestible carbohydrates into SCFAs. Thus, the altered microbiota explains the ability of the host to extract energy from the diet and further stored in the adipose tissue (Turnbaugh et al. 2006; Jumpertz et al. 2011). In support of this notion, we identified several SCFA producing bacteria significantly under-represented or depleted in obese individuals, indicating that SCFAs beneficially regulate host energy metabolism. For example, the relative abundance of some members of *Oscillospira* have been reported to be markedly greater in healthy individuals and associates with human leanness (Beaumont et al. 2016; Konikoff and Gophna 2016; Gophna, Konikoff, and Nielsen 2017). *Oscillospira* utilizes host glycans to produce SCFAs (Konikoff and Gophna 2016; Gophna, Konikoff, and Nielsen 2017) including butyrate, with beneficial effects on insulin sensitivity, body weight control and inflammation (M.-H. Kim et al. 2020). One of the strongest links that has been corroborated across several populations between a gut microbial taxa and BMI involves members of the *Christensenella* genus. They are known to produce SCFAs, acetate and butyrate (Morotomi, Nagai, and Watanabe 2012) and associate negatively with markers of obesity, much in agreement with our findings indicating that *Christensenella* may be important for promoting leanness. We also detected several butyrate producing ASVs including *Eubacterium*, *Alistipes*, *Clostridium* and *Odoribacter* to be proportionally enriched in individuals who were non-obese.

Although we observe more SCFA producing taxa in the non-obese group, we also identify taxa that are SCFA producers in the obese group. Notably we observe that obese individuals presented a greater abundance of *Lachnospira*, a finding consistent with our prior study in the same population (Dugas, Bernabé, et al. 2018), and others (Lippert et al. 2017; Meehan and Beiko 2014; de la Cuesta-Zuluaga, Corrales-Agudelo, et al. 2018). Contrary to our results, other studies have shown that a reduction in the abundance of *Lachnospira* positively associates with obesity (Companys et al. 2021; Stanislawski et al. 2017). It is well known that there are many SCFA producing gut bacteria, raising questions about whether the observed features can be precisely attributed to this mechanism or pathway. However, our predicted functional analysis revealed that genes in the KEGG pathway related to SCFA butyrate synthesis (butanoate metabolism) were significantly depleted or underrepresented in the obese group compared to the non-obese counterparts, which further supports the concept that SCFAs are beneficial. Further, we identified several predicted genes involved in butyrate synthesis via the more-dominant pyruvate pathway in the non-obese group. Altogether, these results suggest that butyrate-producing bacteria may offer protection against obesity (X. Chen and Devaraj 2018). Indeed, butyrate exhibits immunomodulatory effects, improves colon mucosal barrier function, and lowers inflammation.

The SCFA producing microbes dominant in the non-obese group coincided with elevated fecal SCFA levels in these individuals compared with the obese group, which is in line with previous results from other studies that have explored the relation between concentrations of fecal SCFAs and obesity (Yin et al. 2022; Dugas, Bernabé, et al. 2018). Indeed, SCFA supplementation has been documented to protect against a high-fat diet-induced obesity in mice (H. V. Lin et al. 2012; Lu et al. 2016) as well as weight gain in humans (Chambers et al. 2015). Conversely, other studies, mostly from western populations have reported results contrary to our study (Schwiertz et al. 2010; Fernandes et al. 2014; Riva et al. 2017; de la Cuesta-Zuluaga, Mueller, et al. 2018). For instance, de la Cuesta-Zuluaga et al observed associations between elevated fecal SCFA levels, central obesity, gut permeability, and hypertension in a Colombian cohort. The specific mechanisms that explain the higher fecal SCFA levels among obese individuals remain a matter of debate and one hypothesis is that disruptions in the obese gut microbiota may lead to less efficient SCFA absorption, hence the observed increased SCFA excretion (de la Cuesta-Zuluaga, Mueller, et al. 2018). Along the same line of notion, our findings of a negative association between obesity and SCFAs could be related to the consumption of diets enriched in fibers and other dietary precursors of SFCAs resulting in elevated SCFA production compared with SCFA absorption, thereby reducing energy harvesting and its associated storage as fat. Indeed, diets high in fiber and Mediterranean diets correlate positively with weight loss (Hu et al. 2013; Esposito et al. 2011) and increased levels of fecal SCFAs (De Filippis et al. 2016) in human studies. Other possible explanations for the observed divergences between our studies and others might be attributed to differences in population, medication usage, sample size, microbial production capacity and intestinal absorption, underscoring the complex relationships between gut microbiota with SCFA production and host adiposity. Nevertheless, our results demonstrate that the negative associations between obesity and fecal SCFA levels in our study cohort are consistent with the positive associations found between decreased obesity and SCFA synthesizing microbes, although we are aware that fecal SCFA concentrations are not a direct measure of intestinal SCFA production but rather reflect a net result of the difference between SCFA production and absorption (Canfora, Jocken, and Blaak 2015). A measurement of the dynamics of SCFA production and availability with stable isotopes could be determined in future studies. Altogether, the observed differences in SCFA concentrations between obese and non-obese individuals and the several SCFA-producing microbes further reinforce the theory that gut microbiota and its associated SCFA metabolites may have a role in body weight regulation.

Another mechanism by which gut microbiota may contribute to obesity is via the metabolic endotoxemia pathway. Perturbations in the gut microbiota community composition lead to increased production of plasma lipopolysaccharide (LPS) derived from the cell wall of Gram-negative bacteria, provoking low-grade inflammation and increased intestinal permeability which drives adiposity (Zhao 2013; Cani et al. 2008). An increased relative abundance of one ASV assigned to the genus *Dialister* in the gut community of obese individuals was identified from this study. Zhang and colleagues reported proportional increases in *Dialister* in obese persons and suggested that could serve as a potential predictive marker for obesity (Zhang et al. 2021). Additionally, in our recent study (Fei et al. 2021) we observed an increased relative abundance of *Dialister* in subjects with short sleep duration, a condition associated with a chronic inflammatory state. Indeed, *Dialister* has been demonstrated *to* trigger or aggravate host inflammatory response and insulin resistance by releasing more lipopolysaccharides (Yang et al. 2022). To strengthen these findings, we further observed that several genes in the LPS biosynthesis pathway were differentially enriched within the obese group from our predicted functional analysis. Similar findings have previously been reported where the obese microbiota is enriched by LPS metabolism, initiating inflammation-dependent processes associated with the onset of obesity and insulin resistance (Boulangé et al. 2016) and other related metabolic diseases (Yan et al. 2021; Karlsson et al. 2012; Fei and Zhao 2013; Cani et al. 2007; Fei et al. 2021). Collectively, our results demonstrate that obese individuals harbor a marked inflammatory state favoring the development of obesity, and this is in concordance with the associated metabolic endotoxemia pathway linking gut bacteria to obesity.

This study additionally detected marked depletion in pathways involved in cell structure biosynthesis, vitamin B6 biosynthesis, NAD biosynthesis, amino acid metabolism and SCFA synthesis in our predicted metagenome analysis. Thus, our results further suggest that metabolic pathways important for growth, energy homeostasis and the maintenance of normal gut function are disrupted in individuals with obesity. Conversely, in the obese group, we noted an enrichment of formaldehyde assimilation I (serine pathway) pathway. Ubiquitous formaldehyde can be derived from food, the environment and generated endogenously as a result of human and microbial cellular metabolism of many methylated compounds. Endogenous formaldehyde produced at sufficient levels has carcinogenic properties and detrimental effects on genome stability. To counteract this reactive molecule, organisms have evolved a detoxification system that converts formaldehyde to formate, a less reactive molecule that can be used for nucleotide biosynthesis (Reingruber and Pontel 2018; N. H. Chen et al. 2016). Thus, we may infer that the pattern of increased formaldehyde assimilation pathway in our data might result from a defect or diminished capacity of formaldehyde detoxification system pathway, an assumption which requires further verification. A study reported increases in the abundance of formaldehyde assimilation pathway in a depressed group when compared with non-depressed controls (S.-Y. Kim et al. 2022). We are the first to show that the gut of obese participants is enriched in the formaldehyde assimilation pathway. Although we do not understand the mechanistic details, it is known that toxic formaldehyde is generated along with reactive oxygen species during inflammatory processes (N. H. Chen et al. 2016). Thus, an increased capacity for formaldehyde pathway may indicate a microbiome-induced increase in reactive oxygen species in the gut of obese individuals. Indeed, prior work has identified induction of oxygen stress by microbial perturbations as one of the mechanisms by which the microbiome can promote weight gain and insulin resistance (J. Qin et al. 2012). The specific alterations of the gut microbiota and the associated predicted functionality may constitute a potential avenue for the development of microbiome-based therapeutics to treat obesity and/or to promote and sustain weight loss.

## Study strengths and limitations

While our study has several strengths including a large sample size, diverse population along an epidemiological transition gradient with a comprehensive dataset that allowed the exclusion of the potential effects of origin as well as control of potential interpersonal covariates, and use of validated and standard tools for data collection, we acknowledge some limitations as well. First, the cross-sectional nature of our study design is unable to establish temporality or identify mechanisms by which the gut microbiome may causally influence the observed associations. In that regard, we expect that prospective data from the METS cohort study will provide the basis to assess the longitudinal association between gut microbiota composition, metabolites, and obesity, and we have an ongoing study exploring the potential correlations longitudinally. The use of 16S rRNA sequencing in our analysis for inferences on microbial functional ecology inherently has its limitations for drawing conclusions on species and strain level functionality due to its low resolution. Nevertheless, our results provide insight into the relationship between obesity, gut microbiota, and metabolic pathways in individuals of African-origin across different geographies, stimulating further examination of large-scale studies using multi-omic approaches with deeper taxonomic and functional resolution and animal transplantation studies to investigate potentially novel microbial strains and to explore the clinical relevance of the observed metabolic differences.

## Conclusion

This study examined the relationship between the gut microbiota composition and functional patterns, concentrations of fecal short chain fatty acids (SCFAs) and obesity in a large population cohort of African origin, from Ghana, South Africa, Jamaica, Seychelles, and the United States of America, spanning the epidemiologic transition. The Ghanaian cohort exhibited the greatest gut microbiota diversity and the American cohort the least, with corresponding enrichment or depletion in taxa and predicted functional traits. Ghanaian participants were enriched in VANISH taxa reflecting their traditional lifestyle. Significant differences in gut microbiota composition and function were identified in obese individuals compared to the non-obese counterparts. Non-obese individuals were enriched in SCFA-producing microbes which coincided with increased concentration of total SCFA in feces, extending the evidence that SCFAs mediate body weight regulation. The predictive accuracy of the microbiota for obesity status was greatest in low-income countries, and was reduced in high income countries, suggesting that lifestyle traits in high income countries may result in elevated obesity risk even for lean individuals. The specific alterations of the gut microbiota and the associated predicted metabolic function may constitute a potential avenue to guide the development of microbiome-based solutions to treat obesity and/or to promote and sustain weight loss. Thus, further examination of large-scale studies using multi-omic approaches with deeper taxonomic and functional resolution and animal transplantation studies are warranted to confirm the identified taxonomic and metabolic signatures.

## Acknowledgements

We thank the METS participants who continue their ongoing participation in the METS studies, as well as the site-specific clinic staff in Ghana, South Africa, Jamaica, Seychelles and the US. The UCSD Microbiome Core performed sample extractions and library preparation utilizing protocols and primers published on the Earth Microbiome Project website (https://earthmicrobiome.org/). This publication includes data generated at the UC San Diego IGM Genomics Center utilizing an Illumina NovaSeq 6000 that was purchased with funding from a National Institutes of Health SIG grant (#S10 OD026929).

## Contributions

LRD and BTL conceived the study. LRD, CC-K, PB, KB-A, JP-R, TEF, EVL, DR and AL collected human samples and metadata. GEM, CC-K, DR and AL curated metadata. SD performed sequencing of samples. GE-M, CC-K and MGM conducted formal analysis and visualization. JAG and LRD supervised and provided feedback on formal analysis and visualization. GEM, CC-K, MGM, LRD and JAG wrote the original manuscript. LRD secured the funding. All authors edited and approved the final manuscript.

