## Supplemental Files for "Gut microbiota and fecal short chain fatty acids differ with adiposity and country of origin: The METS-Microbiome Study"

*Supplementary Files*

**Supplementary table 1. Alpha diversity between obese and non-obese groups. Alpha diversity estimated by Shannon, Observed ASVs and Faith's PD (Phylogenetic Diversity) between obese and non-obese in each country. Asterisks indicate statistical significance **$p$ **< 0.05 (FDR-corrected). FDR= False Discovery Rate**

| | Observed ASVs | Shannon | Faith's PD |
|---|---|---|---|
| **Country** | **P** | **P** | **P** |
| **Ghana** | 0.0439 * | 0.268 | 0.0393 * |
| **Jamaica** | 0.338 | 0.494 | 0.239 |
| **South Africa** | 0.0488 * | 0.0686 | 0.12 |
| **Seychelles** | 0.206 | 0.752 | 0.186 |
| **US** | 0.759 | 0.246 | 0.901 |

**Supplementary table 2. Gut microbiota (Weighted UniFrac Distance). Adjusted multivariable analysis in the entire cohort (overall) and by each country. Statistical significance from permutational multivariate analysis of variance (PERMANOVA) test, $p < 0.05$. All** ***p*****-values are generated based on 999 permutations**

| | Overall | | Ghana | | South Africa | | Jamaica | | Seychelles | | US | |
|---|---|---|---|---|---|---|---|---|---|---|---|---|
| | $R^2$ | P | $R^2$ | P | $R^2$ | P | $R^2$ | P | $R^2$ | P | $R^2$ | P |
| **Obese** | 0.001 | 0.031 | 0.004 | 0.208 | 0.003 | 0.415 | 0.002 | 0.732 | 0.001 | 0.903 | 0.007 | 0.043 |
| **Sex** | 0.003 | 0.001 | 0.002 | 0.549 | 0.004 | 0.19 | 0.009 | 0.009 | 0.023 | 0.001 | 0.011 | 0.004 |
| **Age** | 0.002 | 0.001 | 0.101 | 0.815 | 0.08 | 0.737 | 0.102 | 0.062 | 0.053 | 0.894 | 0.098 | 0.283 |
| **Country** | 0.118 | 0.001 | | | | | | | | | | |

**Supplementary table 3 – Description of study participants by microbial endotypes. Data are presented as median (interquartile range) for continuous variables, percentages (%) for categorical data. Statistical significance, *p* < 0.05. BMI = Body Mass Index, SCFA = short chain fatty acids**

| | **Bacteroides type** | **Prevotella type** | **p-value** | **q-value** |
|---|---|---|---|---|
| **N** | 820 | 866 | | |
| **BMI** | 29(25,35) | 28(23,34) | 0.003 | 0.004 |
| **Country** | | | <0.001 | <0.001 |
| Ghana | 61 (7.4%) | 265(31%) | | |
| Jamaica | 243(30%) | 117(14%) | | |
| Seychelles | 149(18%) | 176(20%) | | |
| South Africa | 142(17%) | 232(27%) | | |
| US | 225(27%) | 76(8.8%) | | |
| **Sex** | 564(69%) | 503(58%) | <0.001 | <0.001 |
| **Age** | 44(38,50) | 42(36,48) | 0.032 | 0.032 |
| **Obese** | 370(45%) | 346(40%) | <0.001 | <0.001 |
| **Total SCFA** | 3547(2325,4891) | 5300(3591,7672) | <0.001 | <0.001 |
| **Bacteroides** | 1299(315,7521) | 92(18,526) | <0.001 | <0.001 |
| **Prevotella** | 98(25,319) | 1246(198,10382) | <0.001 | <0.001 |

Median(IQR), n (%)

p-value: Wilcoxon rank run test, Pearson's Chi-squared test

q-value: False discovery rate correction multiple testing

**Supplementary table 4. Weight adjusted fecal SCFA levels by country**

| WOMEN | | | | | |
|---|---|---|---|---|---|
| | **Ghana** **n=254** | **South Africa** **n=228** | **Jamaica** **n=263** | **Seychelles** **n=196** | **US** **n=213** |
| Propionate (ug/g) | 30.4 ± 18.0 | 11.6 ± 6.9 | 11.5 ± 8.5 | 14.5 ± 8.5 | 9.1 ± 6.0 |
| Butyrate (ug/g) | 22.1 ± 11.8 | 10.6 ± 6.6 | 8.5 ± 6.8 | 7.0 ± 5.4 | 9.0 ± 6.5 |
| Acetate (ug/g) | 61.3 ± 22.8 | 15.6 ± 7.4 | 24.4 ± 13.3 | 40.9 ± 25.4 | 16.8 ± 11.1 |
| Total SCFA (ug/g) | 115.5 ± 45.0 | 39.5 ± 19.0 | 46.0 ± 26.3 | 63.8 ± 35.1 | 36.4 ± 21.5 |

| MEN | | | | | |
|---|---|---|---|---|---|
| | **Ghana** **n=117** | **South Africa** **n=171** | **Jamaica** **n=133** | **Seychelles** **n=164** | **US** **n=107** |
| Propionate (ug/g) | 34.3 ± 15.9 | 18.6 ± 13.1 | 18.3 ± 13.2 | 21.0 ± 13.9 | 13.1 ± 9.9 |
| Butyrate (ug/g) | 23.1 ± 14.2 | 14.5 ± 9.0 | 13.2 ± 10.1 | 8.6 ± 4.7 | 11.3 ± 6.8 |
| Acetate (ug/g) | 68.9 ± 22.5 | 22.1 ± 9.3 | 27.0 ± 13.3 | 42.2 ± 17.2 | 22.3 ± 13.7 |
| Total SCFA (ug/g) | 128.0 ± 44.2 | 57.3 ± 27.9 | 60.3 ± 30.1 | 72.8 ± 32.0 | 48.4 ± 27.3 |

**Supplementary table 5. Total fecal SCFA by adiposity status; non-obese vs. obese**

| NON-OBESE | | | | | |
|---|---|---|---|---|---|
| | **Ghana** | **South Africa** | **Jamaica** | **Seychelles** | **US** |
| | **n=254** | **n=228** | **n=263** | **n=196** | **n=213** |
| **Propionate (ug/g)** | 34.1 ± 18.3 | 16.8 ± 11.8 | 16.6 ± 13.0 | 19.2 ± 13.2 | 12.7 ± 9.6 |
| **Butyrate (ug/g)** | 23.8 ± 13.2 | 14.3 ± 8.6 | 12.2 ± 9.7 | 8.4 ± 5.9 | 11.1 ± 7.4 |
| **Acetate (ug/g)** | 69.3 ± 22.4 | 21.4 ± 9.2 | 28.3 ± 15.3 | 45.9 ± 24.1 | 23.1 ± 14.2 |
| **Total scfa (ug/g)** | 129.1 ± 45.0 | 54.6 ± 26.3 | 59.0 ± 32.6 | 74.8 ± 37.6 | 48.9 ± 28.4 |

| OBESE | | | | | |
|---|---|---|---|---|---|
| | **Ghana** | **South Africa** | **Jamaica** | **Seychelles** | **US** |
| | **n=89** | **n=141** | **n=178** | **n=132** | **n=201** |
| **Propionate (ug/g)** | 24.9 ± 12.4 | 10.7 ± 6.3 | 10.3 ± 5.9 | 14.6 ± 8.1 | 9.1 ± 6.1 |
| **Butyrate (ug/g)** | 18.7 ± 10.2 | 8.7 ± 5.0 | 7.7 ± 5.6 | 6.6 ± 3.3 | 9.0 ± 6.1 |
| **Acetate (ug/g)** | 48.7 ± 16.9 | 13.2 ± 4.9 | 21.6 ± 9.5 | 34.9 ± 16.8 | 16.0 ± 10.1 |
| **Total scfa (ug/g)** | 93.9 ± 33.9 | 34.1 ± 14.5 | 40.9 ± 18.0 | 57.1 ± 24.0 | 35.4 ± 19.9 |

**Supplementary figure 1**. Differentially abundant taxa among countries (**A-D**) and obese group (**E**) adjusted for BMI, age, sex and country using ANCOM-BC. Bars represent ANCOM-BC estimated log fold change between compared groups (effect size) and error bars, with the 95% confidence interval. ANCOM-BC data for country, representative ASVs with log fold change >1.4 in at least one group are shown. FDR-adjusted (p < 0.05) effect sizes are indicated by *, ** and ***, corresponding to p < 0.05, <0.01 and <0.001 respectively. FDR= False Discovery Rate

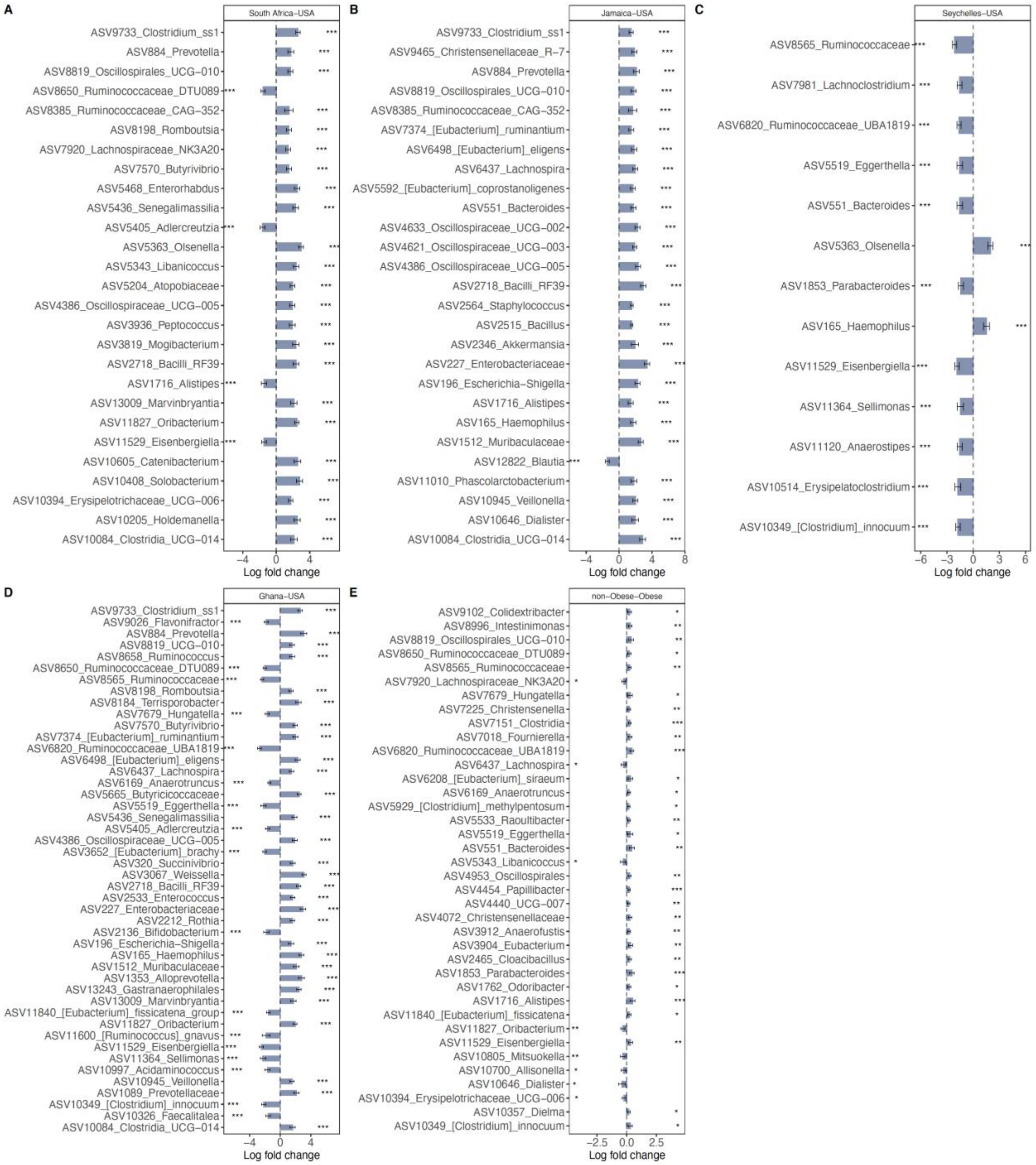

**Supplementary figure 2.** Heatmap representations of the 30 most predictive microbial features (in rows) identified by Random Forest analysis for classification of samples into the various countries (**A**) and into obese and non-obese groups (**B**).

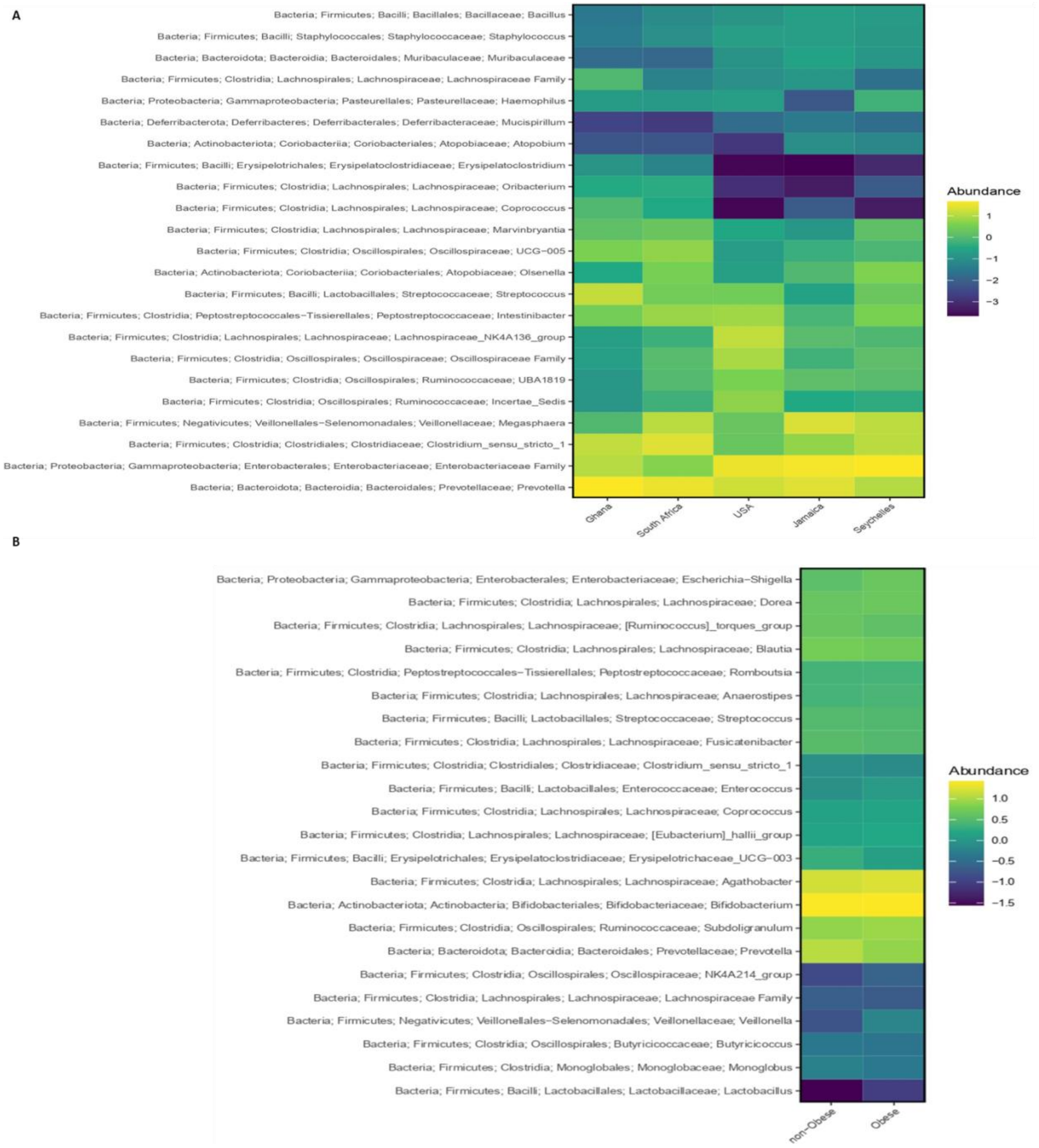

**Supplementary figure 3.** Receiver operating characteristic curves showing the classification accuracy of gut microbiota in a Random Forest model. Classification accuracy for estimating obesity status in (**A**). Ghana; (**B**) South Africa, (**C**). Jamaica; (**D**). Seychelles; (**E**). US are presented. AUC= area under the curve

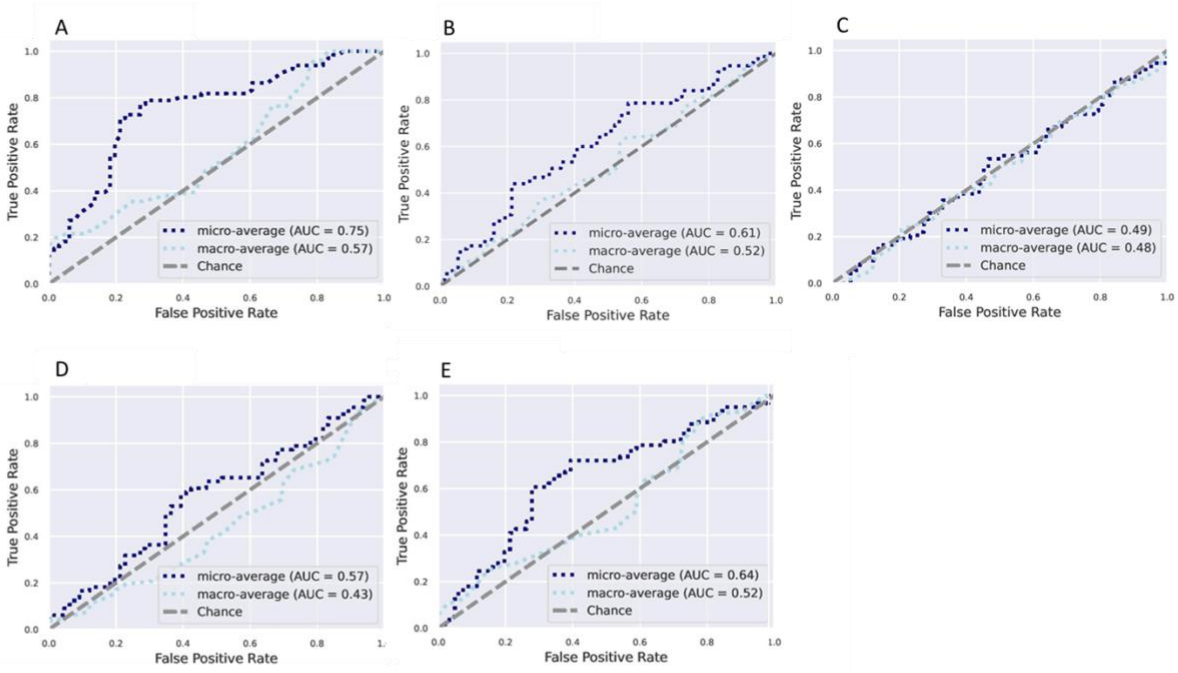

**Supplementary figure 4.** Differentially abundant predicted PICRUSt2 MetaCyc pathways among countries (**A**) and obese group (**B**) adjusted for BMI, age, sex and country using ANCOM-BC. Bars represent the ANCOM-BC estimated log fold change between compared groups (effect size) and error bars, with the 95% confidence interval. ANCOM-BC data for country, representative predicted pathways with log fold change >1.4 in at least one group are shown. FDR-adjusted (p < 0.05) effect sizes are indicated by *, ** and ***, corresponding to *p* < 0.05, <0.01 and <0.001 respectively. FDR= False Discovery Rate

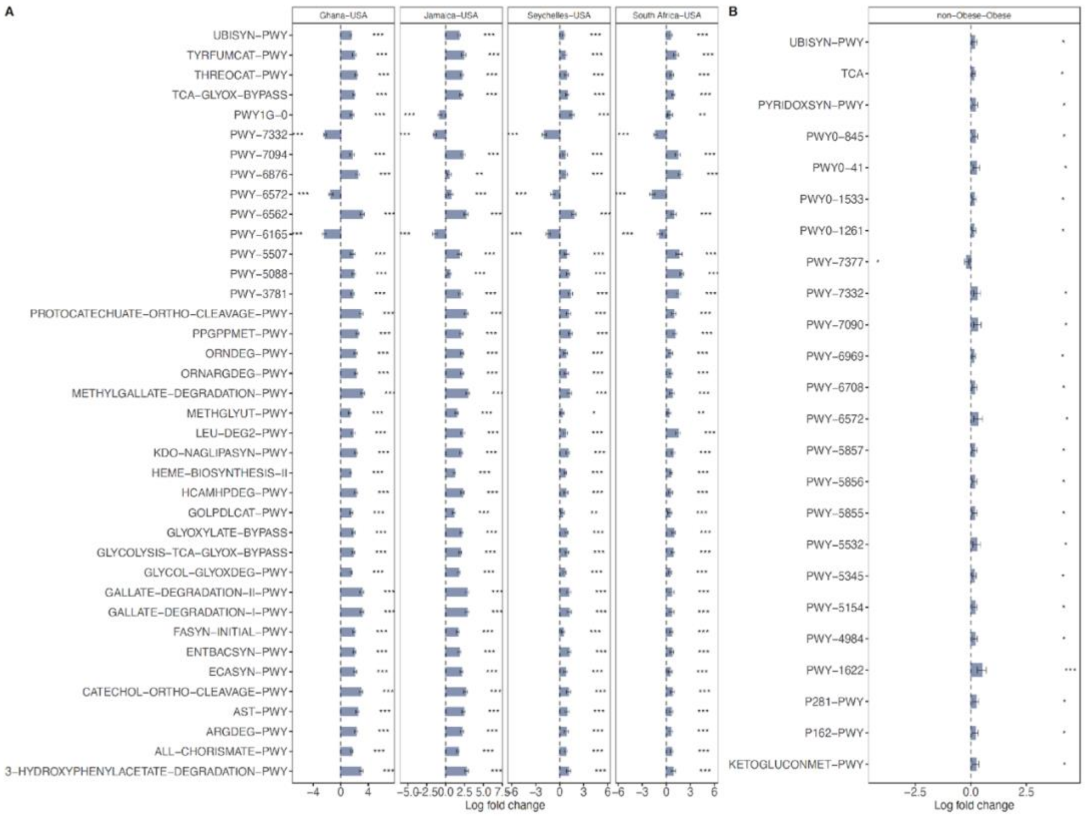

**Supplementary figure 5.** Differentially abundant predicted PICRUSt2 KEGG orthology (KO) in butanoate metabolism pathway among countries (**A**) and obese group (**B**) adjusted for BMI, age, sex and country using ANCOM-BC. Bars represent the ANCOM-BC estimated log fold change between compared groups (effect size) and error bars, with the 95% confidence interval. ANCOM-BC data for country, representative predicted pathways with log fold change >1.4 in at least one group are shown. FDR-adjusted (p < 0.05) effect sizes are indicated by *, ** and ***, corresponding to $p < 0.05$, <0.01 and <0.001 respectively. FDR= False Discovery Rate

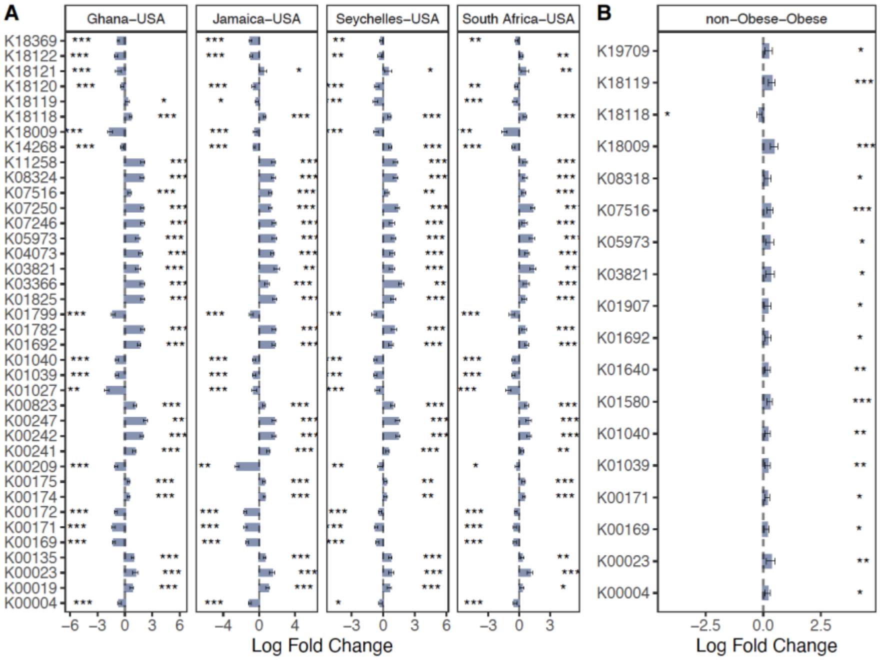

**Supplementary figure 6.** Differentially abundant predicted PICRUSt2 KEGG orthology (KO) in LPS biosynthesis pathway among countries (**A**) and obese group (**B**) adjusted for BMI, age, sex and country using ANCOM-BC. Bars represent the ANCOM-BC estimated log fold change between compared groups (effect size) and error bars, with the 95% confidence interval. ANCOM-BC data for country, representative predicted pathways with log fold change >1.4 in at least one group are shown. FDR-adjusted (p < 0.05) effect sizes are indicated by *, ** and ***, corresponding to $p < 0.05$, <0.01 and <0.001 respectively. FDR= False Discovery Rate

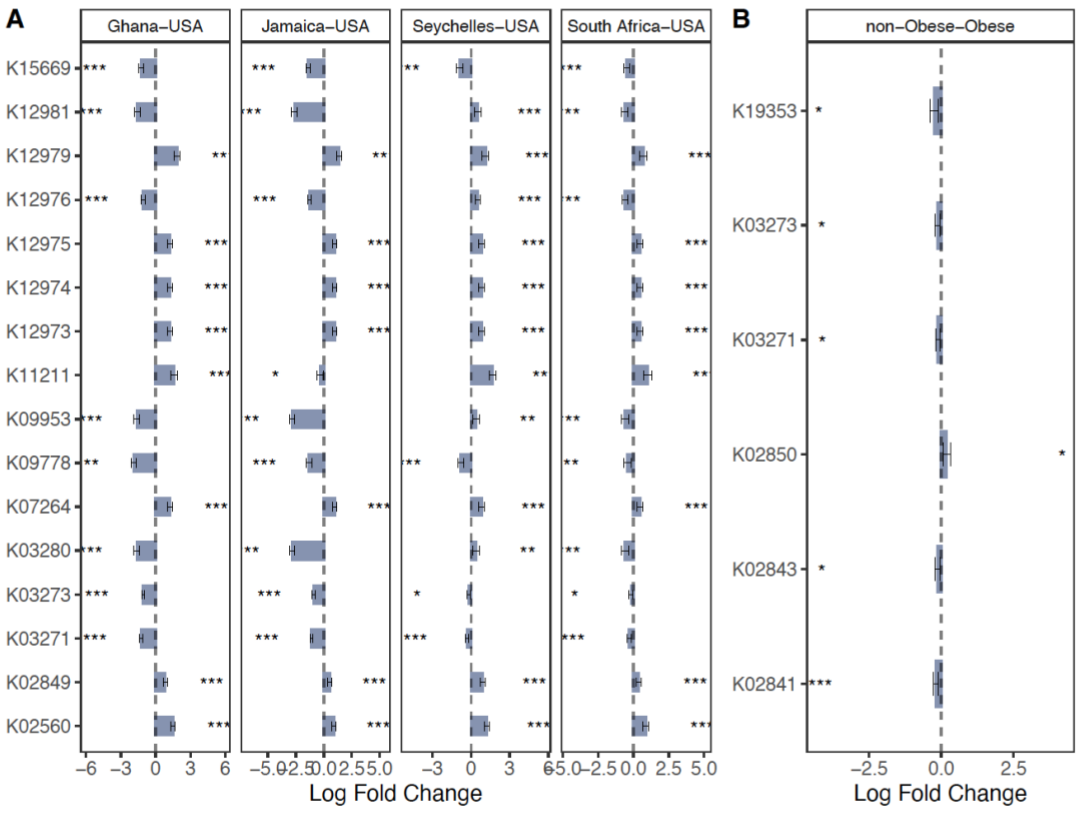
